# Shared neural geometries for bilingual semantic representations in human hippocampal neurons

**DOI:** 10.1101/2025.11.16.688726

**Authors:** Xinyuan Yan, Ana G. Chavez, Melissa Franch, Kalman A. Katlowitz, Ivy Gautam, Brian Kim, Aaditya Krishna, Aadit Shrivastava, Katie Van Arsdel, James Belanger, Assia Chericoni, Taha Ismail, Elizabeth A. Mickiewicz, Danika Paulo, Hanlin Zhu, Alica M. Goldman, Vaishnav Krishnan, Atul Maheshwari, Eleonora Bartoli, Nicole R. Provenza, Seng Bum Michael Yoo, Benjamin Y. Hayden, Sameer A. Sheth

**Affiliations:** Neurosurgery, Baylor College of Medicine; Neurology, Baylor College of Medicine; Neuroengineering Initiative and Electrical and Computer Engineering, Rice University; Department of Bioengineering, Rice University; Department of Biomedical Engineering and Intelligent Precision Healthcare Convergence, Sungkyunkwan University

**Author notes:** These authors contributed equally (Benjamin Y. Hayden, Sameer A. Sheth).

## Abstract

The human brain has the remarkable ability to comprehend and express similar concepts in multiple languages. To understand how it does so, we examined responses of hippocampal neurons during passive listening, directed speaking, and spontaneous conversation, in both English and Spanish, in a small group of balanced bilinguals. We find a small number of putative *cross-language neurons*, whose responses to equivalent words (e.g., “*tierra*” and “*earth*”) are correlated. However, neurons’ semantic tunings differed substantially by language, suggesting language-specific neural implementations. Instead, the crucial driver of translation was a preserved geometric organization of neural responses between the two languages, one that did not depend on neuron level functional overlap. Indeed, that geometry was implemented by a common set of neurons along distinct readout axes; this difference in readout may help prevent cross-language interference. Together, these results suggest that hippocampus encodes a language-independent internal model for meaning.

## INTRODUCTION

Humans have the remarkable capacity to understand and express the same thoughts in multiple languages without confusing them (Bialystok, 2017). For example, in Arnhem Land (Northern Australia), speakers commonly manage six or seven languages (Evans, 2011). A growing body of evidence suggests that bilingual speakers use at least partially shared neural substrates for each of their languages. For example, in fMRI with listening or reading, bilingual brains show broadly overlapping language-network responses across languages (Abutalebi & Green, 2007; Brignoni-Pérez et al., 2022; Cao et al., 2013; de Varda et al., 2025; Klein et al., 1995; Liu & Cao, 2016; Liu et al., 2010; Malik-Moraleda et al., 2022; Sulpizio et al., 2020; Tan et al., 2003). Even constructed languages (e.g., Klingon and Na’vi) reliably recruit the same brain areas as natural languages (Malik-Moraleda et al., 2025). Regions that elicit overlapping responses include classic language regions, such as the inferior frontal gyrus (IFG) and posterior temporal cortex, as well as several others not as closely associated with language. However, these findings do not tell us *how* the brain matches corresponding concepts between languages while simultaneously maintaining a functional separation.

We hypothesized that the bilingual brain uses shared neural geometry for representing meaning in both languages. That is, the brain may have similar sets of coding distances between all pairs of words in high-dimensional neural space. Thus, if “*cat*” and “*dog*” elicit similar neural activation patterns and “*door*” is dissimilar, then “*gato*” and “*perro*” should evoke similar neural activation patterns, and “*puerta*” will be dissimilar. Indeed, this is the strategy used by multilingual LLMs, such as mBERT (Artetxe & Schwenk, 2019; Pires et al., 2019).

A small body of neuroimaging work suggests that the brain may employ shared semantic geometry. In particular, a recent neuroimaging study shows that voxelwise encoding models trained to predict neural activity from language-model embeddings (mBERT) in one language can transfer across languages within the same bilingual participants (Chen et al., 2024). Other studies show cross-linguistic generalization across different monolingual groups listening to translations of the same story (de Varda et al., 2025; Dehghani et al., 2017; Zada et al., 2025), so that encoding models trained on neural responses from speakers of one language predict neural responses in the other (de Varda et al., 2025; Dehghani et al., 2017; Zada et al., 2025). One important study used electrocorticography (ECoG), which provides dense sampling of local field potentials (LFPs), with corresponding bilingual speech in one bilingual patient (Silva et al., 2024). This study found shared articulatory codes sufficient for cross-language decoding, although it did not investigate semantics. Moreover, none of these studies directly examined whether the *shared semantic geometry* (i.e., representational geometry of semantics) exists.

If shared semantic geometry provides the scaffolding that links language to meaning, an important next question is how different languages access this common structure. The properties of representational geometry suggest that rotations of neural axes preserve pairwise relationships, leaving the overall geometric structure invariant (Kriegeskorte & Kievit, 2013; Kriegeskorte & Wei, 2021). As a result, the bilingual brain could access the same semantic geometry through language-specific readout axes. In motor cortex, rotation of neural activity planes enables the same population to produce diverse reaching movements (Sabatini & Kaufman, 2024). Motivated by that work, we reasoned that an analogous process may occur in language.

We are particularly interested in the hippocampus, which has a clear role in encoding word meanings (Binder et al., 2009; Brown-Schmidt et al., 2001; Chavez et al., 2025; Dijksterhuis et al., 2024; Duff & Brown-Schmidt, 2012; Franch et al., 2025; Karkowski et al., 2025; Katlowitz et al., 2025; Piai et al., 2016; Van De Ven et al., 2020). That role in turn is likely part of a broader role in flexibly representing and linking concepts (Bakermans et al., 2025; Bakker et al., 2008; Bernardi et al., 2020; Courellis et al., 2024; Leutgeb et al., 2005; Quiroga, 2012; Yassa & Stark, 2011). The hippocampus has often been ignored in language research, in part due to difficulty imaging it; however, our laboratory has demonstrated that single neurons in the hippocampus track meanings of words during both listening and monolingual speech production (Chavez et al., 2025; Franch et al., 2025; Katlowitz et al., 2025A and B; Mickiewicz et al., 2025; Zhu et al., 2026). Together, these findings suggest that hippocampal neurons form a high-dimensional *semantic geometry* in which the distances between neural activity patterns correspond to the relationships between word meanings.

We recorded single-unit activity from the hippocampus in a rare sample of four fully bilingual (English/Spanish) patients with treatment-resistant epilepsy. Using high-density microelectrodes (N=3) and a Neuropixels probe (N=1), we recorded single units across three studies: (1) passive listening to matched stories in English and Spanish (∼120 minutes each), (2) reading and speaking matched phrases in English and Spanish (using the task developed by Silva et al., 2024), and (3) unconstrained, naturalistic conversations with English and Spanish conversation partners, respectively. Overall, we find evidence of a *shared semantic geometry* that is common to both listening and speaking and that is preserved across the three tasks. At the same time, individual neurons’ semantic tuning curves differed substantially between English and Spanish, and, moreover, each language constructs its semantic geometry through distinct readout configurations of the same neurons. Together, these results clarify how bilingual brains represent shared meanings and suggest the brain uses a readout-based approach to avoiding interference.

## RESULTS

We recorded responses of single neurons in the hippocampus of four bilingual individuals undergoing neurosurgical procedures (**Figure 1A**). All patients acquired both English and Spanish in early childhood (ages 4-5), used both regularly, and were fully balanced bilinguals. Bilingualism was also confirmed informally through conversations in both English and Spanish with both caregivers fluent in each language and by family members. Patients 1-3 underwent stereo-electroencephalography (sEEG) for seizure detection in the epilepsy monitoring unit with an implant strategy that included microelectrodes (AdTech Behnke-Fried) in the hippocampus. Patient 4 underwent anterior temporal lobectomy for medically refractory epilepsy under general anesthesia. Following resection of the neocortex, we recorded from the exposed hippocampus using a Neuropixels probe prior to the hippocampal resection.

**Figure 1.**
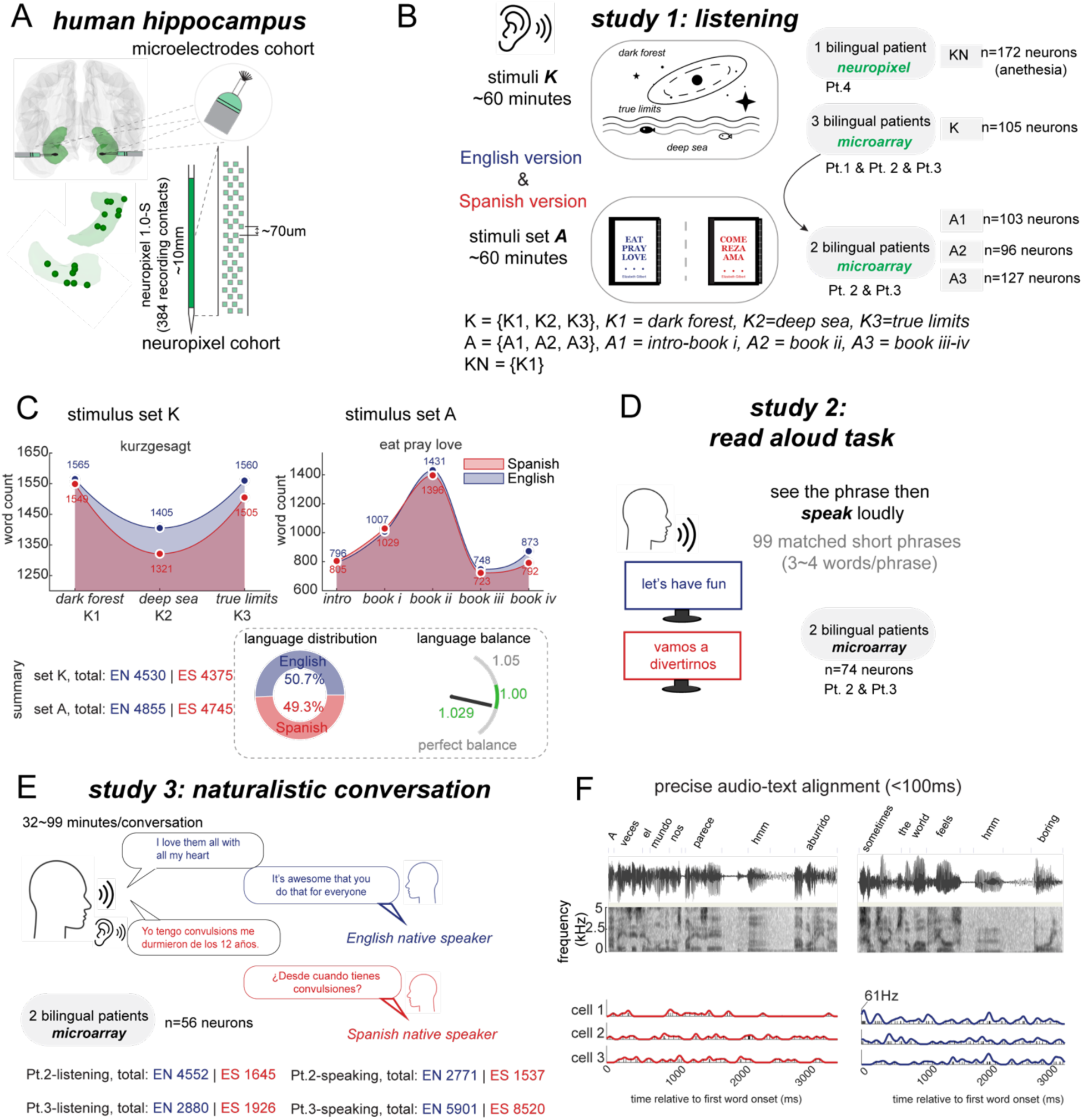
Experimental design for single-neuron recordings during bilingual semantic processing in the human hippocampus. (A) Schematic of recording setup showing bilateral human hippocampus with microelectrode and Neuropixels probe. Green: recording locations. (B) Study 1: the passive listening task involved audio from YouTube videos and an audiobook. Each stimulus set was presented in both English and Spanish versions. (C) Two stimulus sets used for Spanish-English bilingual participants. Word count distributions for stimulus sets K and A showing comparable totals across languages. (D) Study 2: read aloud task. Two patients viewed and spoke aloud 99 matched short phrases (3-4 words per phrase) in both English and Spanish (198 total phrases). (E) Study 3: naturalistic conversation task (32-99 minutes). Two bilingual patients (Patients 2 and 3) engaged in unconstrained conversations lasting 60-90 minutes per session with English and Spanish native speakers. (F) Precise audio-text alignment. Top panel: audio waveform of sample English/Spanish texts. Middle panel: corresponding spectrogram (0-6 kHz) with word boundaries; timestamps are accurate to within 100 ms relative to consensus manual annotations. Bottom panel: example single-neuron responses from three simultaneously recorded cells showing firing rates. Time axis shows milliseconds relative to first word onset for both English (right, blue) and Spanish (left, red) text. Abbreviation: Pt. = Patient.

All patients performed a passive listening task with equivalent speech in English and Spanish (**Figure 1B**, 603 neurons; Stimulus set *K*: 4530 English words, 4375 Spanish words; Stimulus set *A*: 4855 English words, 4745 Spanish words, **Figure 1C**). Two patients (patients 2 and 3) also performed a speech production task in both languages (74 neurons total) in which they saw 99 matching phrases in English and Spanish (198 phrases total) on a computer screen and had to say those words out loud (same task and stimuli as Silva et al., 2024; **Figure 1D**). These two patients also performed a conversation task (Chavez et al., 2025) twice, once in each language, although no effort was made to ensure continuity of topic (**Figure 1E**). Specifically, these two participants took part in conversations in English (durations: 51.37 and 45.5 minutes) and in Spanish (32.42 and 99.98 minutes). Word-level, time-aligned transcripts of the listening and speaking content were produced by combining automated transcription with manual correction in Praat (Boersma, 2001) by trained lab members (**Figure 1F**, see **Methods**).

### A small number of cross-language neurons

We first asked whether neurons show similar responses to co-translated words across language. We initially tried cross-language word-by-word alignment using two open-source software packages (GIZA++ and fast_align) but found their alignments to be insufficiently accurate. We therefore performed a manual English-Spanish alignment (**Figure 2A**). We excluded from analysis words without corresponding meanings and, in many cases, made difficult judgment calls. For example, part of the English text in Stimulus set A reads “*When you’re travelling in India…*” and is translated as “*Al viajar por India…*” In this context, “*When”* and “*Al*” have similar, but not identical, meanings, and were aligned (that is, treated as co-translated pairs). In contrast, “*you’re*” does not have an equivalent word in the Spanish translation and therefore was excluded from the analysis. Meanwhile, “*travelling*” and “*viajar*” have corresponding meanings but different conjugations; given our focus here on semantics over grammar, they were included.

**Figure 2.**
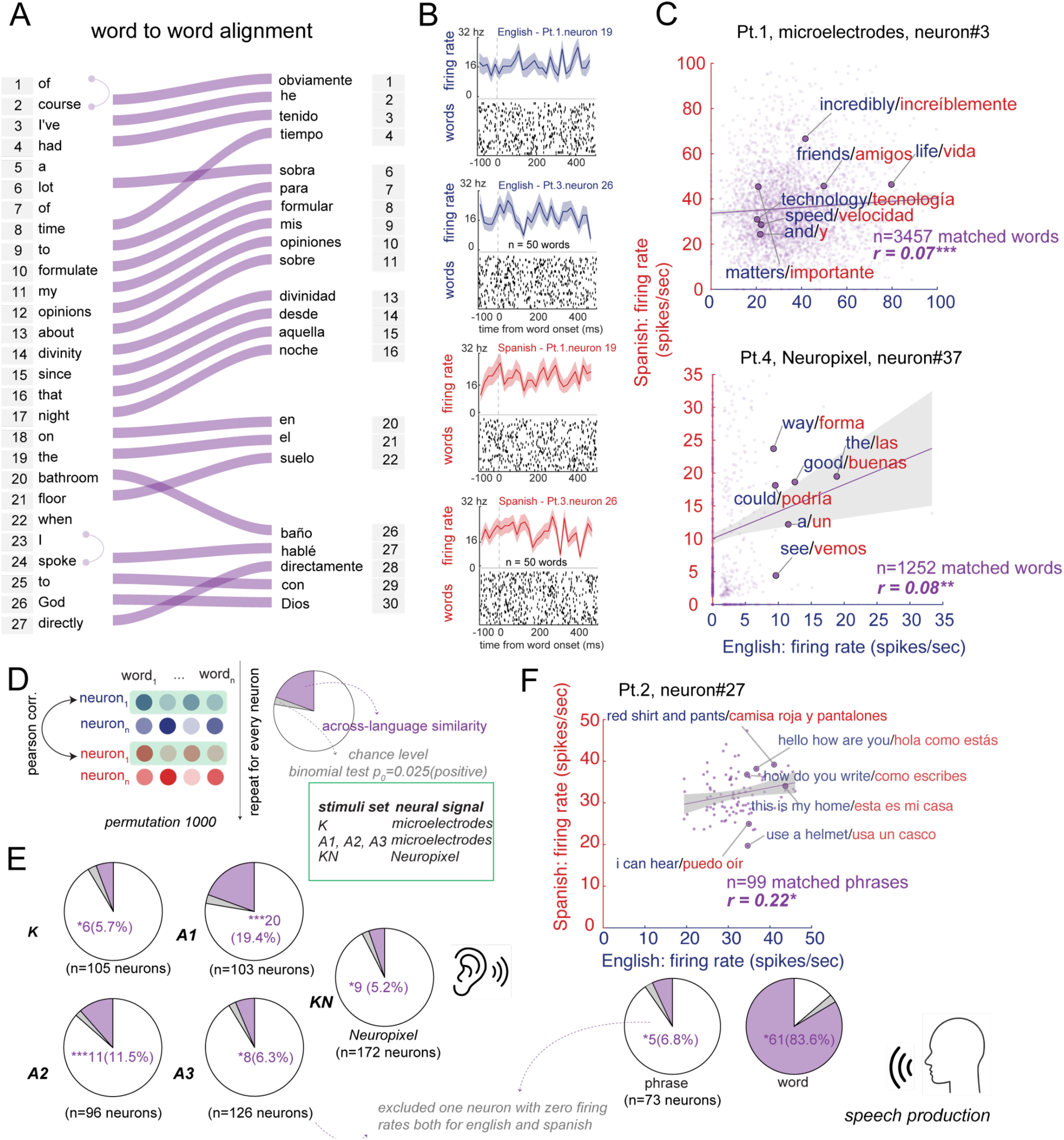
Cross-language single-neuron firing-rate similarity across matched items in passive listening and speech production. (A) Word-to-word alignment mapping between English (left) and Spanish (right) translations. Purple lines connect semantically equivalent words across languages. Numbers indicate word order in each language. Only matched items enter the scatter/correlation analyses below. (B) Peristimulus time histograms (PSTHs) from example neurons. For each panel, top: average response (firing rate, Hz) to 50 matched words in English (blue traces) and Spanish (red traces), bottom: Raster plots below each trace show word-wise spikes. (C) Pearson correlation analysis of firing rates for matched word pairs. Top: Pt. 1 neuron #3 (microelectrodes) showing strong correlation between English and Spanish firing rates for 3457 matched words (r = 0.07, p < 0.001). Bottom: Pt. 4 neuron #37 (Neuropixels) displaying similar language-invariant responses. Each dot is one matched word pair, x-axis is the unit’s firing rate in English, y-axis represents the unit’s firing rate in Spanish. For visualization, several items are annotated (e.g., *friends/amigos*, *life/vida*). Shaded regions: +/- SEM. (D) For each neuron, we computed the Pearson correlation between firing rates across translation-matched English-Spanish word pairs and assessed significance with a one-tailed permutation test (positive direction; 1000 shuffles of Spanish indices). We then counted neurons with r>0 and two-tailed permutation p<0.05 and tested whether this fraction exceeds chance using a right-tailed binomial test with (p₀ = 0.025). (E) A set of “cross-language neurons” across all stimulus conditions. Pie charts show the proportion of neurons exhibiting significant cross-language similarity for each stimulus set (purple), with grey portions indicating the expected 2.5% false positive rate under the null hypothesis of no cross-language positive relationships. (F) A set of “cross-language neurons” during speech production (Study 2, read aloud task). Example neuron from Pt. 2 shows correlated firing patterns when patient was speaking matched English-Spanish phrase pairs. Shaded regions: +/- SEM. (***p<0.001, **p<0.01, *p<0.05). See also Table S1, Table S2 and Table S4.

Example responses (**Figure 2B**) illustrate neurons that show comparable evoked responses to co-translated words across languages. Example units showed significant positive relationships across thousands of matched items, indicating that words that elicit above-average (e.g., “*friends*”/“*amigos*”) or below-average (e.g., “*and*”/“*y*”) firing rates in English tend to do so in Spanish as well (**Figure 2C**). To systematically identify putative *cross-language neurons*, we computed a Pearson correlation across all matched words for every neuron (e.g., for stimulus set K, n=3457 matched words for Patients 1-3; n=1252 matched words for Patient 4) and used a binomial test to determine whether the proportion of significant correlations exceeded chance levels (**Figure 2D**, **Methods**). A small number of neurons with zero firing rates across all words in either language was excluded (n=1 in dataset A3, listening task; n=1 in read aloud task).

A subset of neurons met the cross-language neuron criterion for both listening (**Figure 2E**) and speaking (**Figure 2F**; also see **Table S1-Table S2**). Note that sessions of the listening task (Study 1) and speech production (Study 2) were collected on different days, so we cannot tell if the same neurons contribute to speaking and listening. This difference between phrase level and word level likely reflects signal attenuation from averaging across words within each phrase, as phrase-level effect sizes were substantially smaller than word-level effect sizes (mean r = 0.03 vs. 0.29; Wilcoxon signed-rank test, p<0.001). All phrase-level cross-language neurons were also identified at the word level, and 67.2% of word-level-only cross-language neurons showed phrase-level correlations above their permutation null (binomial test, p = 0.006), confirming that the cross-language signal exists at the phrase level but maybe too weak for per-neuron detection. We found essentially no evidence for “anti-cross-language” neurons (i.e., neurons with significant negative correlations between neural responses for matched words across languages). The lack of *a priori* equally likely anti-cross-language neurons in all datasets supports the idea that cross-language neurons, while rare, are not simply a statistical artifact. However, the overlap in firing rate magnitude does not necessarily imply that tuning is identical across languages, nor does it imply that they play a crucial role in translation.

### English-Spanish tuning profiles rarely align at the single-neuron level

Individual neurons respond systematically to word meanings with *semantic tuning functions* (Chavez et al., 2025; Franch et al., 2025; Jamali et al., 2024). To test how semantic tuning functions relate across languages, we first extracted contextual word embeddings using multilingual BERT (mBERT, **Figure 3A**, **Methods**), a transformer model that learns cross-linguistic representations through masked language modeling (Devlin et al., 2019).

**Figure 3.**
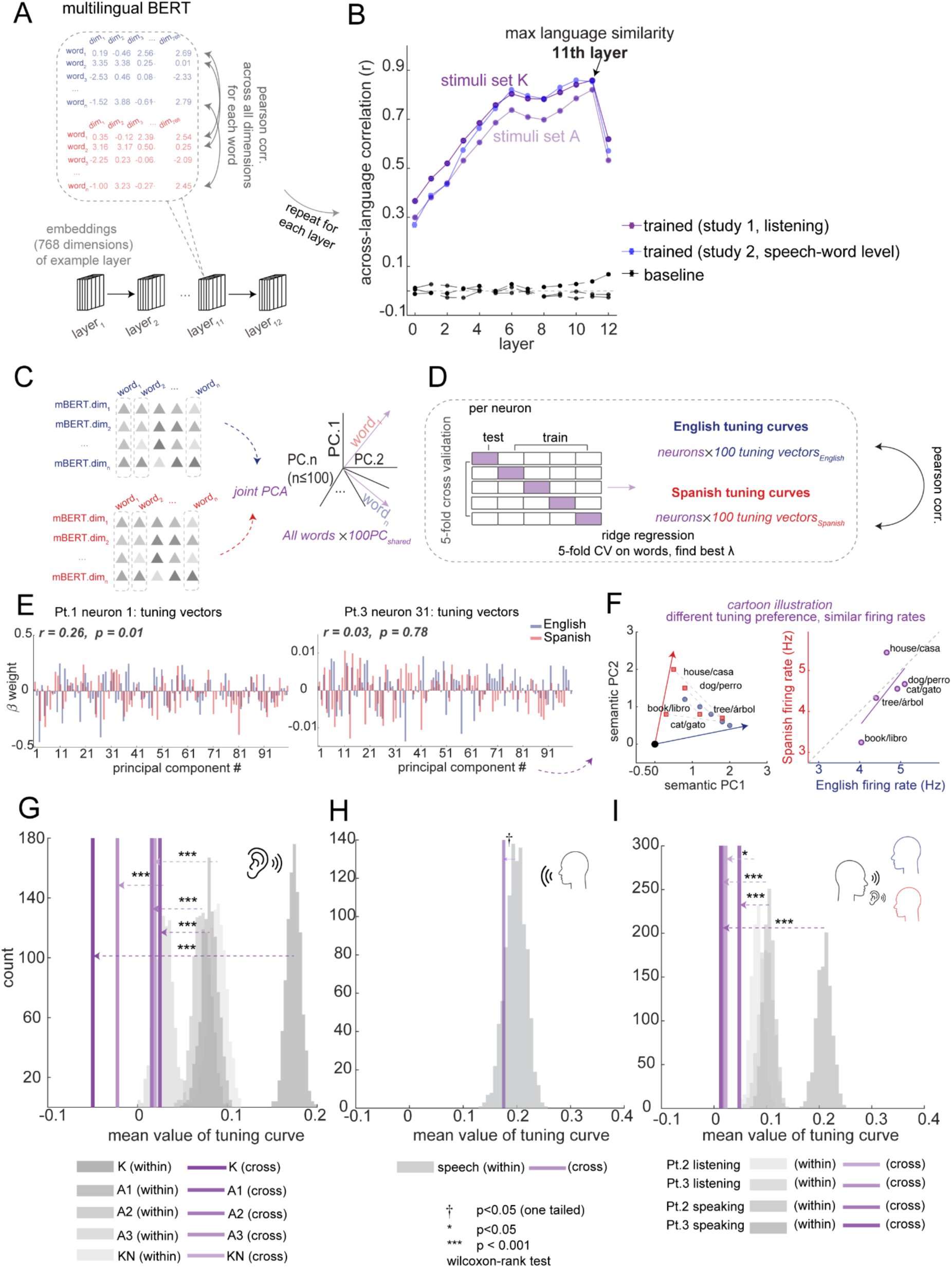
Semantic embeddings and tuning curves similarity at a single neuron level. (A) Multilingual BERT (mBERT) architecture and contextual embedding extraction pipeline. (B) Cross-language correlation (purple curves; two stimulus sets: K = *Kurzgesagt* podcasts; A = audiobook; blue curves, words from read aloud task) rises with depth and peaks at layer 11, which we therefore used in all analyses in passive listening (Study 1), speech production (Study 2) and conversation (Study 3). (C) Fitting tuning curves in a shared semantic coordinate system. English and Spanish words’ embeddings from each dataset are concatenated and projected onto a joint PCA, yielding a 100-PC shared basis. (D) For each neuron and each language separately, we fit ridge regression from the 100 PCs of all words to firing rates of all words, selecting *λ* by 5-fold cross-validation. The resulting beta is the neuron’s semantic tuning curve for that language (for each neuron, there are 100 *β* (i.e., 100 tuning vectors) for English, and 100 *β* for Spanish). Across-language tuning curve similarity of each neuron was defined by the Pearson correlation between English and Spanish tuning curves. (E) Different tuning preferences across languages in the example neurons. Bars show β weights across the 100 shared PCs for two representative neurons (left panel from example “cross-language” neuron, right panel from example “non-cross-language” neuron). Neuron-wise English-Spanish tuning curve correlations are not significant. (F) Cartoon illustration to explain different tuning preferences across languages but may result in similar firing rates. Left: English and Spanish use different readout directions in the semantic PC1-PC2 plane; Right: projections of the *same words* onto those directions are nevertheless similar, yielding similar firing rates. This visualizes “output-similar does not mean recipe-identical”: firing rates similarity across languages can arise from different weights of semantic components. (G, H, I) Mean value of across-language tuning curves similarity (purple vertical ticks) of all neurons compared with within-language tuning curves similarity in (G) passive listening (Study1), (H) speech production (Study2), and (I) naturalistic conversation (Study3). Histograms show the distribution of within-language tuning curve similarity. Across all tasks, the across-language tuning curve similarities are below the within-language similarity, demonstrating that tuning curves were not well aligned at the single-neuron level. (***p<0.001, *p<0.05, †p<0.05 (marginally significant if two-tailed, significant if one-tailed)) See also Figure S1.

For stimulus set A, we combined excerpts from the audiobook presented across three recording sessions (A1, A2, A3). Sentences were tokenized using mBERT preserving their original narrative sequence (**Methods**). For every matched translation pair (e.g., *“dog”/“perro”*), we calculated the correlation between English and Spanish word embeddings (768-dimensional vectors from mBERT itself, not neural data) at each mBERT layer and computed the average correlation across words. Layer 11 (out of 13) yielded the highest cross-language similarity (**Figure 3B**). This result is consistent with reports that deeper layers carry semantic information and yield stronger cross-lingual alignment (e.g., Mousi et al., 2024). We therefore used Layer 11 embeddings for Study 1 and Study 3. For Study 2 (read aloud task), words were produced in short phrases without narrative context. We therefore extracted mBERT embeddings for each phrase independently (**Methods**). Both Layer 10 and Layer 11 showed the highest cross-language similarity for matched translation pairs (**Figure 3B**, blue line); we used Layer 11 for consistency with Studies 1 and 3.

We projected both languages’ embeddings into a common space by concatenating English and Spanish embeddings and applying joint PCA, retaining 100 principal components (**Figure 3C**). This approach ensured both languages were projected onto identical semantic axes. To validate this shared PCA space, we conducted two control analyses. First, for each shared component, we computed the fractional contribution of each language to the total variance explained by that component, which yielded nearly balanced contributions (English = 0.509; Spanish = 0.491). Second, we confirmed that each principal component derived from the joint PCA showed a strong correlation between English and Spanish scores (average correlation, r = 0.501, all PCs, p < 0.001), indicating that both languages aligned along the same semantic axes. These results confirm that the shared PCA captured common axes for both languages.

For every neuron, we then fit ridge regressions to map these shared semantic features to firing rates separately for English and Spanish. To avoid overfitting, we selected a single ridge penalty (λ) per neuron via 5-fold cross-validation that minimized the average normalized error across both languages (see **Methods**). A neuron’s cross-language tuning similarity was defined as the Pearson correlation between its English and Spanish tuning profiles (i.e., the tuning vectors across the shared PCA components from ridge regression). Significance was assessed by 1000-fold permutation test (**Figure 3D**).

Significant correlations between English and Spanish tuning vectors were rare, indicating that neurons do not generally have matching overall tuning in the two languages (**Figure 3E, F**; also see **Figure S1**). We next asked whether the divergence in tuning curves between English and Spanish was greater than within-language tuning similarity. For each study (Study1, listening; Study 2: read aloud; Study 3: conversation) separately, we split the English and Spanish words into two halves (independently, 1000 random splits), refit ridge weights on each half, and computed within-language split-half tuning curve correlations per neuron. The mean value cross-language tuning correlation fell at the extreme left of the within-language tuning curve correlation distribution (near the 0.3 percentile in Study 1 and Study 3). Cross-language tuning profile correlations were significantly lower than the within-language tuning profile correlation (Wilcoxon signed-rank test for all stimuli sets in Study 1 and Study 3. All z-values < -2.20, all p <0.02; **Figure 3G, I**). For Study 2 (speech production) at the word level, however, the cross-language tuning correlation did not significantly differ from within-language tuning correlations (Wilcoxon signed-rank test, p = 0.06, **Figure 3H**). This result is consistent with the high proportion of cross-language neurons observed at the word level in Study 2 (83.6%, Figure 2F). Cross-language neurons (n = 115) showed significantly higher cross-language tuning similarity (mean = 0.14, SD = 0.11) than other neurons (n = 560, mean = -0.01, SD = 0.15; Welch’s t-test: t(204) = 11.83, p < 0.001), suggesting that when words are produced in isolation, hippocampal neurons may exhibit largely language-invariant semantic tuning. Taken together, semantic tuning profiles generally differ across languages, with the notable exception of isolated word production, where cross-language neurons maintain similar tuning across English and Spanish.

### Population geometry preserves cross-language semantic structure

We next asked whether *semantic geometry* is shared across English and Spanish. We utilized representational dissimilarity matrices (RDMs), which capture the pairwise distances between word representations and are invariant to axis swaps, rotations, and linear remixing (Diedrichsen & Kriegeskorte, 2017; Kriegeskorte et al., 2006; Kriegeskorte & Wei, 2021). These transformations (e.g., rotation) can change single-neuron tuning functions without altering the population geometry, making RDMs ideal for testing whether English and Spanish maintain the same semantic geometry.

For both languages, we constructed RDMs from populational hippocampal neural activity by concatenating word-evoked firing-rate vectors across neurons and computing pairwise cosine distances (“*neural semantic maps*”) among words. In parallel, we computed mBERT RDMs from mBERT Layer 11 embeddings of the same words (“*LLM semantic maps,*” **Figure 4A-F** and **Methods**). We asked two questions: (1) how similar is the *neural semantic map* to the *LLM semantic map* (brain-model alignment)? (2) How similar is the *neural semantic map* across English and Spanish within the brain (cross-language neural RDM correlation)?

**Figure 4.**
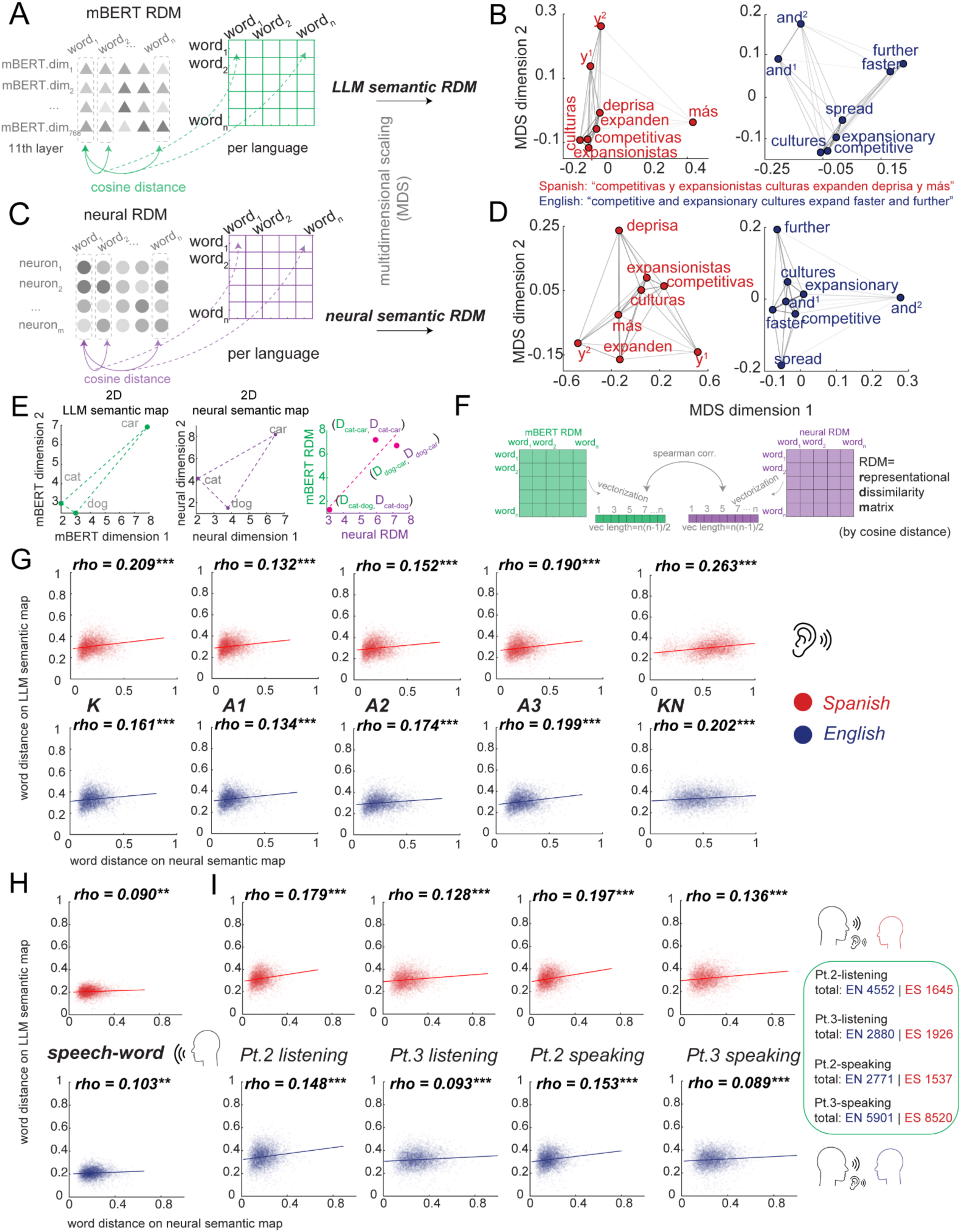
Neuronal population semantic geometry preserved across English and Spanish. (A) Construction of LLM semantic map. Word embeddings from mBERT are used to compute pairwise cosine distance matrices for each language. (B) Multidimensional scaling (MDS) visualization of LLM semantic map for words from the example sentence. Gray lines represent distances between word pairs. Semantic geometry constructed from mBERT embeddings shows that words with close semantic distances in English maintain similar relationships in Spanish. (C) Construction of neural semantic map. Pairwise neural distance matrices are computed from population firing rate vectors (neurons shown as circles) using cosine distance. The resulting matrices capture how the neural population differentiates between words: the more dissimilar two words are, the larger their distance in neural space. (D) Multidimensional scaling (MDS) visualization of neural semantic map for words from the example sentence. Neural population geometry shows that neurons organize words using similar patterns in English and Spanish. (E) Schematic illustrating the relationship between LLM semantic map and neural semantic map analyzed in panels G-I. If “cat-dog” are close in semantic space, they also tend to be close in neural space, appearing near the diagonal line in the rightmost panel. (F) Distance matrix correlation computation. Semantic distance matrices (green, from word embeddings) and neural distance matrices (purple, from population responses) are vectorized and correlated using Spearman correlation. (G) Brain-model alignment in passive listening (Study 1). Scatter plots show correlations between words distances in LLM-mBERT space (x-axis) and neural space (y-axis) for datasets K, A1, A2, A3, and KN-Neuropixel. All datasets show significant positive correlations (r = 0.114-0.237, all p < 0.001). (H) Brain-model alignment in read aloud task (Study 2). Scatter plots show correlations between word distances in LLM-mBERT space (x-axis) and neural space (y-axis) for words in short phrases. The results show significant positive correlations (rho= 0.090 for Spanish, and rho=0.103 for English, all p < 0.01). (I) Brain-model alignment in naturalistic conversation (Study 3). Word distances in neural space aligned to the words distances in LLM-mBERT during listening and speaking. (***p<0.001, **p<0.01) See also Figure S2, Table S3.

We used Spearman tests to analyze representational similarity, following best practices (Nili et al., 2014). Brain-model alignment was significant within both languages. Specifically, across datasets in Study 1, the *neural semantic map* significantly correlated with the *LLM semantic map* (Spearman’s rho = 0.132-0.263, all p < 0.001, permutation test; **Figure 4G**). That is, word pairs that are distant in the *LLM semantic map* (e.g., “*human*” and “*galaxy*”) are also distant in the *neural semantic map*, while close word pairs (e.g., “*cat*” and “*dog*”) in the *LLM semantic map* are close in the *neural semantic map*. These alignments are robust in both English and Spanish. Study 2 (read aloud task) also showed similar brain-model alignment (**Figure 4H**; Spearman’s rho = 0.090-0.103, all p < 0.01, permutation test). Study 3 (conversation) showed similar brain-model alignment, despite using different sets of words: correlations were significant during listening (Spanish: Pt. 2, rho = 0.179, Pt. 3, rho = 0.128; English: Pt. 2, rho = 0.148, Pt. 3, rho = 0.093) and speaking (Spanish: Pt. 2, rho = 0.197, Pt. 3, rho = 0.136; English: Pt. 2, rho = 0.153, Pt. 3, rho = 0.089), all p < 0.001 (**Figure 4I**). Together, these results confirm that the hippocampus and mBERT share semantic geometry both for English and Spanish.

Crucially, the *neural semantic map* was preserved across languages (example see **Figure 5A-B**). We directly compared the neural RDMs of the two languages by correlating pairwise neural distances between words in English with the corresponding pairwise neural distances between their Spanish translations across all matched word pairs. We found significant positive correlations for all passive listening datasets (rho = 0.255-0.477, all p < 0.001; **Figure 5C&D** & **Method**). Cross-language alignment was also observed in Study 2 (rho = 0.247, p < 0.001; **Figure 5E&F**). Permutation controls show that shuffled Spanish word labels produced null distributions centered near zero, with observed correlations falling far outside these distributions for both passive listening and speech production (all z-values>3.0, all p <0.001; **Figure 5D, F**).

**Figure 5.**
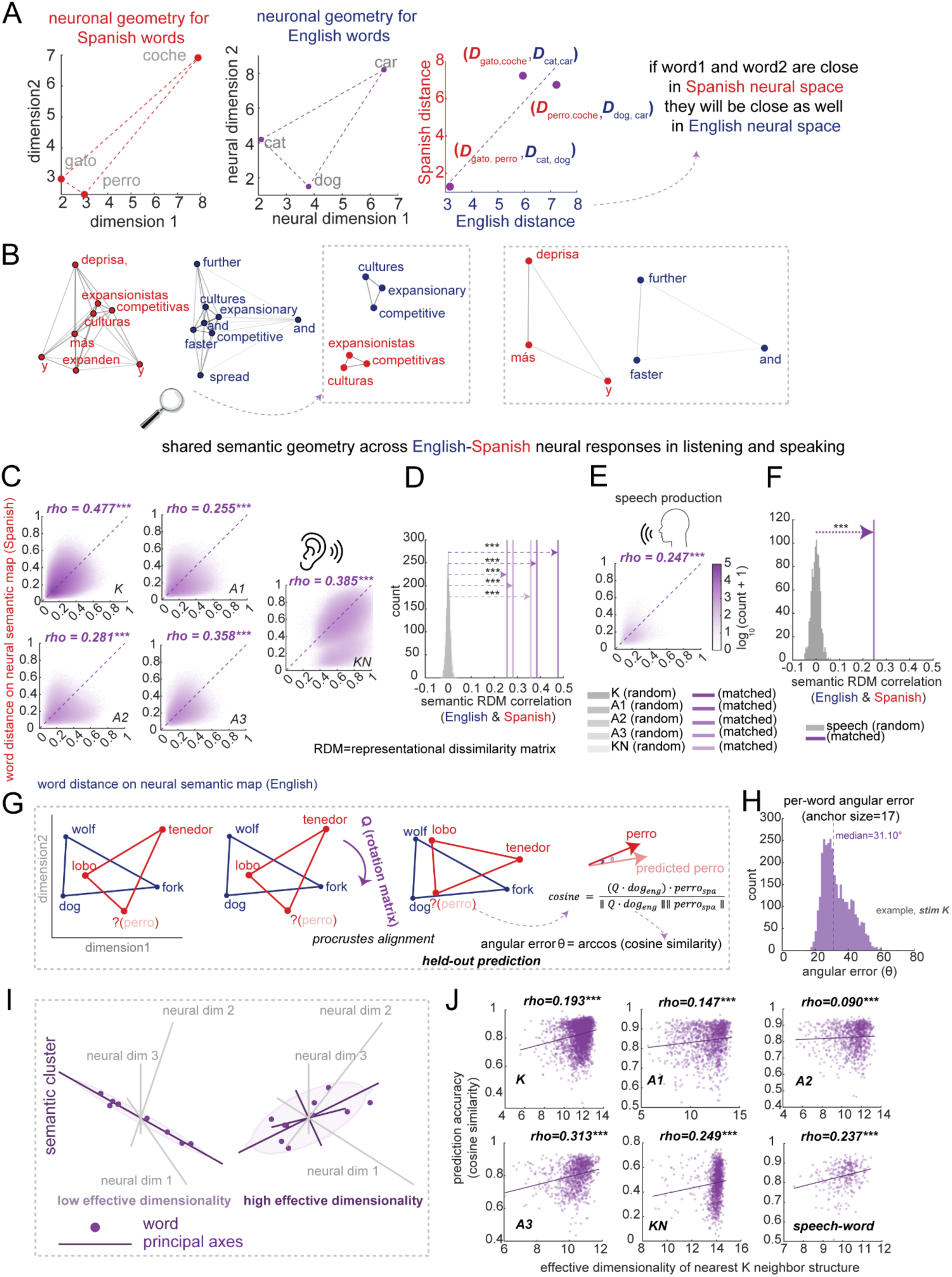
Cross-language alignment of neural population geometry reveals shared semantic geometry. (A) Schematic illustrating cross-language neural distance preservation. Left: Example words in Spanish neural space. Middle: Same words in English neural space. Right: Cross-language distance correlation plot where each point represents a word pair. (B) Zoomed views highlighting semantic clusters within each language from the example sentence in Figure 4. Words with close semantic relationships cluster together in both languages. (C) Shared neural semantic geometry between English and Spanish. The two-dimensional histograms show the joint distribution of pairwise neural distances measured in English (x-axis) and in Spanish (y-axis) for corresponding word pairs. Color intensity represents log_10_(count + 1) of word pairs falling within each bin (60×60 bins, color scale 0-5; **Methods**). Purple gradient from white (sparse) to dark purple (dense) indicates concentration of data points. Diagonal dashed line represents perfect cross-language correspondence. Passive listening datasets (K, A1, A2, A3, KN-Neuropixel) show significant correlations. (D) Permutation test validation for passive listening. Histogram shows null distribution of cross-language neural distance correlations generated by randomly shuffling word-pair assignments (1000 permutations, gray bars). (E) Speech production (read aloud task) shows significant alignment (rho = 0.247, p < 0.001). Dense clustering along diagonal indicates that word pairs maintaining specific distances in English neural space preserve similar distances in Spanish neural space. (F) Permutation test for speech production (read aloud task). Similar analysis showing that the observed cross-language correlation (purple vertical line) significantly exceeds the null distribution from random word-pair assignments (gray histogram), validating semantic specificity of neural geometric alignment during speech. (G) Schematic of local Procrustes prediction method. The aim is to predict the target word (e.g., “perro” in light red) in Spanish given known structure from matched English words. For the translation-equivalent target word in English (e.g., “dog”), we identified its *K* nearest semantic neighbors in English neural space (left, for visualization, only 3 words shown; Methods), learned a rotation matrix Q from the cross-language response pairs of these neighbors (middle), and applied Q to predict the target word’s Spanish neural response (right). The predicted Spanish response (light red) is compared with the true Spanish response vector (dark red) via cosine similarity and angular error θ (arccos(cosine similarity)). (H) Per-word angular error distribution (example from stimulus set *K*). Histogram of angular error (θ = arccos(cosine similarity)) between predicted and actual Spanish response vectors at optimal anchor size (k = 17) for stimulus set K. Median angular error = 31.10°. See **Figure S3E** for all datasets. (I) Effective dimensionality schematic. Low effective dimensionality (left): anchors cluster (neighborhoods cluster) along a single neural dimension, providing limited geometric constraints. High effective dimensionality (right): anchors span multiple neural dimensions. (J) Structured neighborhoods predict better. Relationship between effective dimensionality of anchor sets and prediction accuracy across datasets. Higher dimensionality is associated with better prediction (Spearman ρ = 0.09-0.31, all p < 0.001), indicating that structured neighborhoods outperform redundant ones. (***p<0.001). RDM=representational dissimilarity matrix (by cosine distance) See also Figure S3, Table S3.

We also compared cross-language RDMs derived from mBERT embeddings. Given the strong brain-model alignment observed within each language (**Figure 4G&H**), it is unsurprising that mBERT exhibited similarly high correlations between English and Spanish RDMs (rho = 0.651 for K; rho = 0.620 for KN; rho = 0.504 for A1; rho = 0.467 for A2; rho = 0.529 for A3; all p < 0.001).

### Local geometric structure enables cross-language prediction

Previous studies have used learned encoding weights between word embeddings and neural responses to predict neural activity in another language (Chen et al., 2024; de Varda et al., 2025; Zada et al., 2025). However, none have tested whether such prediction is possible at the single-word level. This question is crucial, because we can often infer the meaning of an unfamiliar word from its context, even without explicit translation knowledge, a form of zero-shot learning (Pourpanah et al., 2023; Socher et al., 2013). For example, in English we know that “*dog*” and “*wolf*” are close in meaning, while both are far from “*fork*”.

To test whether preserved semantic structure could support single-word neural prediction, we used local Procrustes alignment (**Figure 5G**, **Methods**). For each target word, we identified its k nearest neighbors in English neural space (excluding the target itself for held-out prediction aim), learned a rotation matrix from these neighbors’ cross-language response pairs, and predicted the target’s Spanish response. Prediction accuracy was quantified as angular error between predicted and actual vectors. We optimized anchor size (k = 15-19 across datasets; **Figure S3A**) and found median angular errors of 30-61° for listening and word-speech tasks (see example for stimulus set *K* in **Figure 5H**, all results see **Figure S3E**), significantly exceeding null (all p < 0.001). Sliding window analysis showed that even distant neighbors (positions 50-100) predicted nearly as well as the nearest neighbors (<1° decay), indicating a near-global rather than strictly local transformation (**Figure S3B & C**).

We further found that the neighbor-cluster tightness strongly predicted accuracy (Spearman rho= 0.37-0.59; **Figure S3D**). Most importantly, higher effective dimensionality of anchor sets, indicating rich structured rather than redundant neighborhoods (**Figure 5I**), was associated with better prediction (Spearman rho = 0.09-0.31, all p<0.001; **Figure 5J**). This aligns with work showing that high-dimensional neural geometry supports cross-condition generalization (Fusi et al., 2016; Bernardi et al., 2020). Structured neighborhoods spanning multiple neural dimensions provide richer geometric constraints for estimating the rotation, enabling better generalization to held-out words than redundant low-dimensional clusters. Note that Study 3 was not included in cross-language neural comparisons (**Figure 5**) because conversational content differed between languages, precluding word-by-word matching.

### Cross-language neurons are not essential for cross-language neural semantic geometry

We next confirmed that the previously identified cross-language neurons are not responsible for the cross-language geometry match. To test this, we repeated the analyses above excluding the cross-language neurons (that is, the ones that showed similar English-Spanish firing profiles, identified in **Figure 2**, n=115). Even after excluding these neurons, we still observed brain-model alignment within languages (all p<0.001) and cross-language neural semantic map correlations (all p<0.001). Critically, the drop in correlation upon removing cross-language neurons did not exceed what would be expected from removing a random subset of equal size (**Table S3**).

### Shared semantic geometry emerges from language-specific semantic axes within a common neural ensemble

Similar pairwise distances can, in principle, be maintained even if the underlying coordinate axes are altered, leaving the geometry unchanged (Kriegeskorte & Wei, 2021, **Figure 6A**). Moreover, English and Spanish may engage distinct subsets of neurons to construct their respective semantic spaces; or they may recruit the same neuronal population but implement different rotations or re-weightings of the neuronal axes to form language-specific semantic axes. Our results support the latter account.

**Figure 6.**
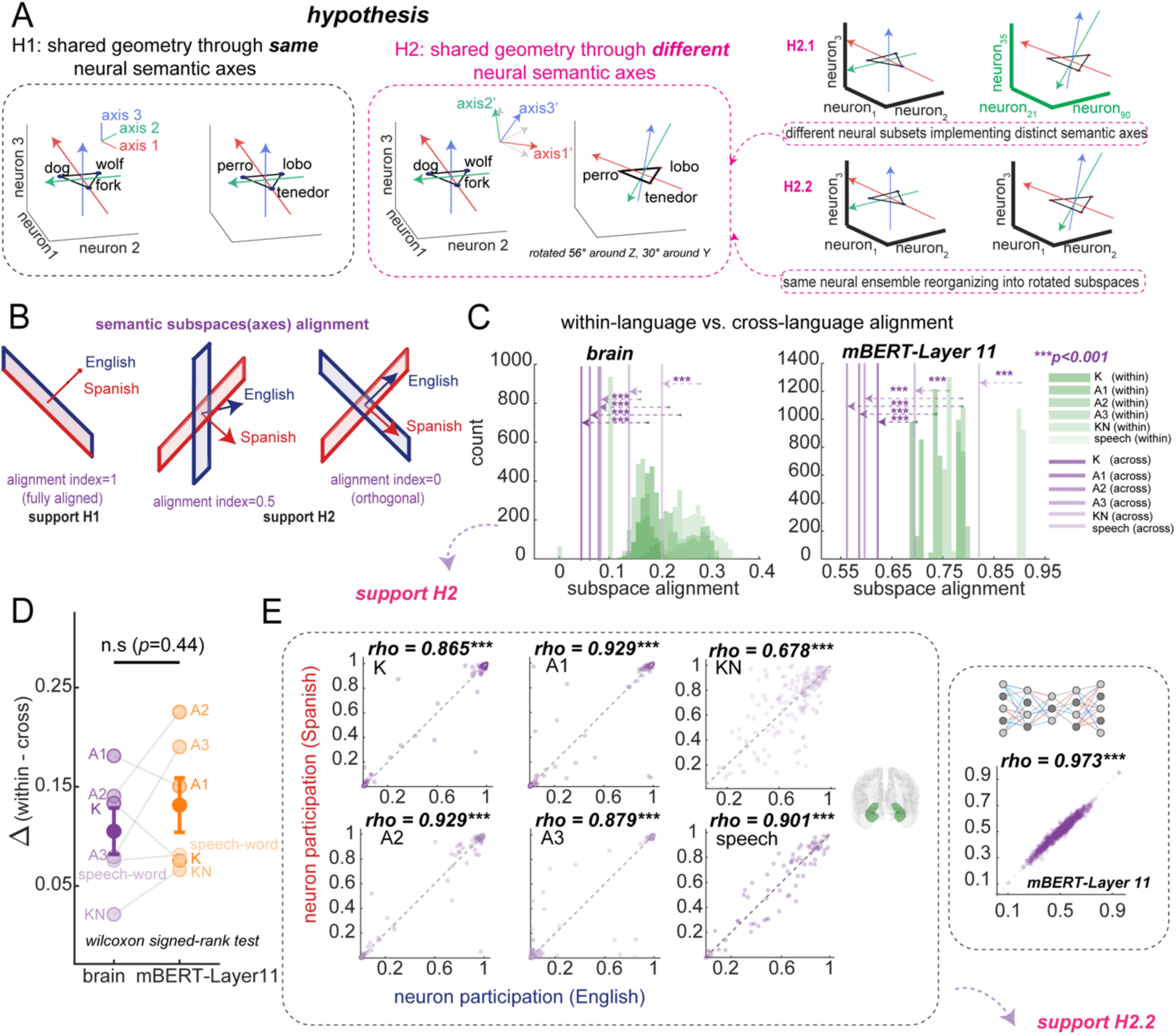
English and Spanish recruit the same group of neurons but different semantic readout axes to support a shared semantic geometry. (A) Conceptual illustration showing that the shared semantic geometry across-language can be preserved in two ways: either by relying on the same semantic axes (left panel, hypothesis *H1*), or by altering the underlying semantic axes (right panel, red and green axes rotated 56° in X-Y plane, blue axes tilted 30°, supporting hypothesis *H2*). If *H2* is correct, then there are two possible mechanisms that could give rise to different neural semantic axes across languages: *H2.1*, English and Spanish may engage different subsets of neurons to construct their semantic spaces, or *H2.2*, they may recruit the same subset of neurons but use rotations or reweighting of the neuronal axes to form language-specific semantic axes. (B) Conceptual illustration of possible subspace alignments between languages. Fully aligned (1.0, support H1), partially aligned (0.5, support H2), and orthogonal (alignment = 0, support H2). (C) Eigendecomposition of the centered RSM (representational similarity matrix, 1-RDM) yields the principal semantic axes (eigenvectors) for each language. Then we build the subspaces based on the top eigenvectors (which captured 95% of variance) and test the across-language alignment (**Methods**). The principal semantic axes for English and Spanish were less aligned compared to within-language subspaces both for mBERT-Layer 11 and hippocampus. (D) The reduction in alignment from within-language to cross-language was comparable between the hippocampus and mBERT (Layer 11). (E) Per-neuron participation strength in the top subspace (explaining > 95 % variance) was highly correlated between languages both for hippocampus and mBERT (Layer 11), showing that both English and Spanish rely on nearly the same group of neurons to form their semantic readout axes, even though their semantic subspaces differ in directions. (***p<0.001). See also Figure S4, Figure S5 and Table S5.

We applied eigendecomposition to the representational similarity matrices (RSMs, 1-RDM, **Figure 6B**) to extract the principal axes (eigenvectors) and quantified cross-language subspace alignment using variance-capture alignment indices (**Figure 6C**, Elsayed et al., 2016; **Methods**). We term the eigenvectors of the semantic similarity matrix as *semantic axes*, representing the principal directions of meaning in population space. These axes can be interpreted as *semantic readout axes*, because they specify linearly decodable directions that downstream circuits could access (Okazawa et al., 2021).

Despite the similar semantic geometry, the manifold constructed from semantic axes for English and Spanish were less aligned than the within-language subspaces both for mBERT (K, A1, A2, A3, KN, speech-word, all p<0.001) and hippocampus (K, A1, A2, A3, speech-word, all p<0.001; KN-Neuropixel, p>0.05, **Figure 6C**). But the absolute across-language subspace alignment value in mBERT is substantially larger than values (both within and across-language subspace alignment) observed in the brain (**Figure S4A**). The reduction in alignment from within-language to cross-language conditions was comparable between the hippocampus and mBERT (hippocampus=0.11±0.06 vs. mBERT-Layer11=0.13±0.07, p = 0.44, Wilcoxon signed-rank test; **Figure 6D**). For both systems, the across-language alignments are significantly higher than randomly shuffled distribution (unmatched words) across all stimulus sets (all p<0.001), indicating that English and Spanish implement similar semantic geometry through differently oriented, partially independent semantic axes.

We then examined the neural coordination structure through eigendecomposition of neuron×neuron covariance matrices, which quantify how neurons co-vary across words within each language (**Methods**). Each neuron’s participation strength (the sum of squared loadings across the top eigenvectors, reflecting how heavily it contributes to these coordination modes), was highly correlated between languages (**Figure 6E**; K: rho=0.87; A1: rho=0.93; A2: rho=0.93; A3: rho=0.88; KN-Neuropixel: rho=0.68; speech-word, rho=0.90, all p<0.001; mBERT results, see **Figure S4B**). These effects are not driven by a small subset of extreme neurons (see control analysis in **Figure S5**).

## DISCUSSION

We examined responses of hippocampal neurons in four bilingual English-Spanish speakers with implanted microelectrodes. We find evidence for a shared cross-linguistic neural embedding space for both listening and speaking for the two languages. That is, words with similar meanings in English and Spanish evoke similar neural patterns of responses across the two languages, leading to shared semantic geometry. Each neuron had distinct semantic tuning for the two languages, and the shared semantic geometry arose from the different semantic readout axes of two languages. Thus, cross-language shared representation of concepts appears to be a complex emergent phenomenon that takes place not primarily through specialized neurons or even a translation axis, but rather on the manifold, through shared geometry (Ebitz & Hayden, 2021; Perich et al., 2025; Vyas et al., 2020). Together, these results suggest a specific solution to the problem of translation, one that gives a central role to shared coactivation patterns in driving the ability to move between languages (Conneau et al., 2017).

While important neuroimaging studies (Chen et al., 2024; Correia et al., 2014; de Varda et al., 2025; Van de Putte et al., 2017; Zada et al., 2025) demonstrate that the brain encodes semantic features in similar way across languages, our single-unit findings take a further step. We quantify the population neuron-level embeddings for each individual word and reveal how every word occupies a specific position within a high-dimensional semantic space. The across-language prediction based on geometric features revealed that the preserved semantic geometry in hippocampal population codes is sufficient to infer the embedding vectors of a hidden word, indicating that shared conceptual geometry is sufficient for translation, and that direct word-to-word mapping is not necessary.

We also consider the relationship between mBERT and the brain. At the population level, the representational geometry of mBERT (layers 9-11; **Figure S2A-S2C**) resembles that of the hippocampus. Both represent words as high-dimensional vectors with linearly decodable structure and exhibit cross-language geometric alignment in which translation equivalents occupy similar relative positions (Devlin et al., 2019; Pires et al., 2019; Wu et al., 2022). Importantly, this alignment is layer-selective: brain-mBERT correlations were negative in early layers, shifted positive around layers 4-6, and peaked in layers 9-11 (**Figure S2**), indicating that the hippocampus aligns specifically with higher semantic layers rather than mBERT’s full hierarchy. Both systems achieve shared geometry through language-specific subspaces constructed from overlapping populations (neurons in the brain, embedding dimensions in mBERT; Figure S4), though mBERT’s cross-language subspace alignment far exceeds that observed in the brain at every layer. At the single-unit level, a fundamental difference emerges: the vast majority of ‘units’ are cross-language correlated units, 100% units of higher layers of mBERT (>=layer 6) are cross-language correlated units (**Table S5**). Meanwhile, neurons with similar tuning functions (“cross-language neurons”) are rare in our dataset (5.2%-19.4%, but 83.6% in speech production only). What might account for this difference? One potential reason is a scale mismatch: mBERT similarities are computed over thousands of dimensions that average across many representational factors, whereas a single neuron may reflect a narrow, idiosyncratic subset of features. A second possibility is that mBERT is partly driven by surface form regularities—shared subword structure, orthography, and distributional cues (for example, “*universe*” and “*universo*”) whereas neurons in semantic regions may be more strictly organized around conceptual content. Finally, there may be a deeper representational difference: mBERT encodings are relatively linear and factorized, so related words align in clean vector subspaces, whereas neural codes may be more curved, multi-lobed, and context-dependent (Perich et al., 2025).

### Cross-language neurons

We identify a new class of what we call “cross-language neurons,” whose responses to co-translated words are correlated. Such neurons would have similar responses to, for example, “*maybe*” and “*quizás*.” These neurons may contribute to translation; however, they do not play an essential, or even a major role: even when we exclude them from our analysis, cross-language semantic geometry is robustly maintained (**Table S3**). Moreover, while their tuning functions are more aligned than other neurons, they are not perfectly aligned, indicating that they are not simply amodal conceptual representations. Consistent with this, their response fields span multiple semantic categories (e.g., **Figure 2C**), which differentiates them from the canonical amodal concept-related neurons, concept cells (Quian Quiroga et al., 2005; Quian Quiroga, 2012). Word-level decomposition confirms that their cross-language correlations are driven by a relatively sparse subset of words. These words are constrained along fewer latent semantic axes in mBERT space than other neurons that did not show cross-language correlation (**Table S1-S2**). We speculate that they may be the extreme end of a larger distribution of neurons whose shared population-level geometry contributes to translation.

### How geometry provides simultaneous shared meaning and language-specific readout

Bilinguals need to resolve a control problem that monolinguals do not face: they must not only select a word that matches their intended meaning but also select between available equivalent words in different languages. That is, a bilingual speaker who wants to order a glass of water may choose a different equivalent term in El Paso and Ciudad Juárez. That may feel effortless, but it requires a specialized representational scheme that allows for simultaneous, readout-dependent semantic coding, while also allowing for translatability. In other words, bilingual representations are a manifestation of a broader category of generalization/differentiation problems. The need for simultaneous generalization and differentiation is one that is difficult to solve with single neurons but that is readily solved with population codes (Barak et al., 2013; Bernardi et al., 2020; Johnston et al., 2024; Libby & Buschman, 2021; Tang et al., 2020; Tian et al., 2024; Xie et al., 2022). We speculate that language context (provided by surrounding words, interlocutor cues, and even general knowledge) could gate which readout axes’ rotation is active (Rigotti et al., 2013).

#### Limitations of the Study

One limitation of our study is the low number of patients that we recorded. The main reason for this low number is the rarity of balanced bilingual patients who acquired both languages early in life with access to single units in hippocampus. However, our core finding of shared semantic geometry was replicated across all listening datasets, extended to speech production (i.e., read-aloud task), and was significant in all four patients individually. Second, we did not test additional languages. While English and Spanish are different, they are genetically related and many words are cognate; it would be valuable to test these ideas on unrelated languages. Third, our data cannot address how the shared semantic geometry is formed. Future studies that track participants as they learn a new language could reveal how semantic geometry emerges and becomes aligned across languages. Likewise, recordings from patients who learned their languages at different life stages could shed light onto how existing languages serve as a scaffold for learning new ones (Odlin, 2003). Finally, some of our data come from an anesthetized patient whose anterior temporal lobe had been resected prior to recording (but the Spanish data from the same patient have never been published or analyzed). Both the anesthesia and the resection may have altered patterns of responding (but see Katlowitz et al., 2025A), although we find largely similar results between this patient and our other three.

## STAR★METHODS

### Ethics approval

The experiment’s procedures were conducted by the standards set by the Declaration of Helsinki and approved by the Institutional Review Board for Baylor College of Medicine, Houston, TX (H-50885 and H-18112). Participants provided written informed consent after the experimental procedure had been fully explained and were reminded of their right to withdraw at any time during the study.

### Patient recruitment

Four fully balanced Spanish-English bilingual epilepsy patients (daily use of both languages since age 4) participate in the study and were aware that participation was voluntary and would not affect their clinical course. Patients’ age ranged from 26-54 years old (mean ± s.e.m = 38.50 ± 7.08 years), with four female patients. None of the patients reported explicit memory of intraoperative events after the case when discussed in the post-operative care unit or while recovering in the hospital the next day.

### Experimental stimuli in study 1 (Figure 1B&1C)

Neural activity was recorded using either Neuropixels probe (P04) or microelectrodes enabling single neuron isolation (P01-P03). During passive listening tasks (Study1), participants were presented with naturalistic audio stimuli in both Spanish and English. To assess the robustness of our findings across different content domains, we employed two distinct stimulus sets (Set A (audiobook: “*eat, pray, love*”) and Set K (*kurzgesagt*)) that varied in their linguistic properties and semantic content. The stimuli consisted of four distinct recordings: three educational podcast episodes from *Kurzgesagt* (K: K1-“dark forest”; K2-“true limits”; K3-“deep sea”) and one complete audiobook (A: audiobook “*Eat, Pray, Love*” by Elizabeth Gilbert). The stimulus presentation varied by participant based on recording availability: P04 (with Neuropixels probe) listened to K1 only (we termed this dataset *KN-Neuropixel*), P01 listened to K1-K3, while P02 and P03 listened to both stimulus sets (K1-K3 and A). Due to clinical care requirements, the audiobook stimulus (A) was recorded across three non-consecutive days for P02 and P03 (Day 1: A1 (books 0-1); Day 2: A2 (book 2); Day 3: A3 (books 3-4)), whereas all podcast stimuli (K1-K3) were collected in single sessions.

### Experimental stimuli in study 2

We adapted the bilingual speech neuroprosthesis paradigm from Silva et al. (2024) to investigate cross-linguistic neural encoding during active speech production (in our study, we called it as “read aloud task”). The experimental design employed a carefully curated vocabulary of bilingual short phrases presented in an isolated-target task paradigm, where participants attempted to produce individual words in either English or Spanish upon visual cue presentation. The core vocabulary comprised 99 matched English and Spanish short phrases (3-4 words; see Supplementary Table 7 of Silva et al., 2024). During this task, participants viewed a single target word displayed on a screen, then the participants attempted to produce the word overtly. The task design incorporated several key features to optimize neural signal quality. First, phrases were presented in randomized order across both languages to prevent anticipatory effects and ensure that neural responses reflected language-specific processing rather than sequential patterns. Second, the visual presentation format remained consistent across all trials, with text displayed in the same font, size, and screen position to minimize visual processing confounds.

### Neuropixels data acquisition setup and intraoperative recordings in the hippocampus

Neuropixels 1.0-S probes (IMEC) with 384 recording channels (total recording contacts = 960, usable recording contacts = 384) were used for recordings (dimensions: 70μm width, 100μm thickness, 10mm length). The Neuropixels probe, consisting of both the recording shank and the headstage, were individually sterilized with ethylene oxide (Bioseal, CA, (Coughlin et al., 2023)). Our intraoperative data acquisition system included a custom-built rig including a PXI chassis affixed with an IMEC/Neuropixels PXIe Acquisition module (PXIe-1071) and National Instruments DAQ (PXI466 6133) for acquiring neuronal signals and any other task-relevant analog/digital signals respectively. Our recording rig was certified by the Biomedical Engineering at Baylor St. Luke’s Medical Center, where the intraoperative recording experiments were conducted. A high-performance computer (10-core processor) was used for neural data acquisition using open-source software such as SpikeGLX 3.0 and OpenEphys version 0.6x for data acquisition (AP band (spiking data)), band-pass filtered from 0.3kHz to 10kHz was acquired at 30kHz sampling rate; LFP band, band-pass filtered from 0.5Hz to 500Hz, was acquired at 2500Hz sampling rate). We used a “short-map” probe channel configuration for recording, selecting the 384 contacts located along the bottom 1/3 of the recording shank.

Audio was played via a separate computer using pre-generated wav files and captured at 30kHz or 1000kHz on the NIDAQ via a coaxial cable splitter that sent the same signal to speakers adjacent to the patient. MATLAB (MathWorks, Inc.; Natick, MA) in conjunction with a LabJack (LabJack U6; Lakewood, CO) was used to generate a continuous TTL pulse whose width was modulated by the current timestamp and recorded on both the neural and audio datafiles. Online synchronization of the AP and LFP files was performed by the OpenEphys recording software. Offline synchronization of the neural and audio data was performed by calculating a scale and offset factor via a linear regression between the time stamps of the reconstructed TTL pulses and confirmed with visual inspection of the aligned traces.

Acute intraoperative recordings were conducted in brain tissue designated for resection based on purely clinical considerations. The probe was positioned using a ROSA ONE Brain (Zimmer Biomet) robotic arm and lowered into the brain 5-6mm from the ependymal surface using an AlphaOmega microdrive. The penetration was monitored via online visualization of the neuronal data and through direct visualization with the operating microscope (Kinevo 900).

Reference and ground signals on the Neuropixels probe were acquired separately by connecting to a sterile microneedle placed in the scalp (separate needles inserted at distinct scalp locations for ground and reference respectively).

#### Motion analysis

The motion-corrected location estimates were obtained at a 250Hz sampling frequency using the DREDge algorithm. This signal was down sampled to 10Hz. The power spectrum of the calculated motion was then estimated using Welch’s overlapped segment averaging estimator for frequencies between 0.1 and 3Hz. The amount of motion was defined as the root mean square error of the location trace of the probes center relative to its average location.

#### Thresholding method

Spike detection was performed using an adaptive thresholding algorithm designed to achieve a target firing rate of approximately 20 Hz per channel. A threshold was set dynamically for each channel based on signal characteristics and noise level. For each channel noise level was estimated using the mean absolute deviation (MAD), calculated as MAD = median (|x|)/0.6745, where x is the band-passed filtered signal. Channels with MAD values below 1×10-6 were classified as dead channels and excluded. An iterative binary search algorithm was used to determine the optimal threshold multiplier for each channel. Different values were tested within a range. Negative-going threshold crossings were considered a neuronal spike. If the iterative firing rate was too high, the threshold was made more negative and vice versa for a low firing rate. A refractory period of 1ms was also used to eliminate some level of noise.

### Microelectrodes recording in the hippocampus

The hippocampus was not a seizure focus area in any of the patients included in the study. Single neuron data were recorded from stereo-EEG (sEEG) probes, specifically Ad-Tech Medical probes in a Behnke-Fried configuration. Each patient had an average of three probes terminating in the left and right hippocampus. Electrode locations were verified by co-registered pre-operative MRI and post-operative CT scans. Each probe includes eight microwires, each with eight contacts, designed explicitly for recording single-neuron activity. Single neuron data were recorded using a 512-channel *Blackrock Microsystems Neuroport* system sampled at 30 kHz. To identify single neuron action potentials, the raw traces were spike sorted using the *Wave_clus* sorting algorithm (Chaure et al., 2018) and then manually evaluated and modified to improve sorting. Noise was removed, and each signal was classified as multi or single unit using several criteria: consistent spike waveforms, waveform shape (slope, amplitude, trough-to-peak), and an exponentially decaying ISI histogram with no ISI shorter than the refractory period (1 ms). The analyses here used all single and multiunit activity.

The initial number of neurons was simply the total number of neurons that we collected. All preprocessing was done by a lab member blind to the goals of the study. In preprocessing (spike sorting), we only kept the isolated neurons. We analyzed all collected neurons, and did not restrict analyses to task-responsive neurons.

### Firing rate responses to words

In Study 1 (listening) and Study 3 (conversation-listening), firing rates were computed from spikes occurring 200 ms after word onset to 200 ms after word offset, accounting for transmission delays to hippocampus (Charest et al., 2009; Chavez et al., 2025). In Study 2 (read aloud task) and Study 3 (conversation-speaking), the analysis window was shifted 200 ms earlier (from 200 ms before word onset to 200 ms before word offset) to capture neural activity during speech preparation (Chavez et al., 2025). Spike counts were divided by window duration and multiplied by 1000 to yield spikes per second.

### Audio transcription

After experiments, the audio.wav file was automatically transcribed using Assembly AI, a state-of-the-art AI model trained to transcribe speech. Speaker diarization was achieved with Assembly AI’s internal speaker diarization model. The transcribed words, corresponding timestamps, and speaker labels output from these models were converted to a TextGrid and then loaded into Praat, a well-established software for speech analysis (https://www.fon.hum.uva.nl/praat/). The original *.wav* file was also loaded into Praat. Trained lab members manually inspected the spectrograms, timestamps, and speaker labels, correcting each word to ensure that onset and offset times were precise and assigned to the correct speaker (this process took about 4-5 hours per minute of speech). The TextGrid output of corrected words and timestamps from Praat was converted to an Excel file and loaded into Python (version 3.11) for further analysis.

### Temporal dynamics of semantic encoding (Study 2)

To characterize the timing of semantic encoding during reading aloud, we performed a sliding window analysis aligned to two events: visual onset (when the phrase appeared on screen) and speech onset (when the participant began speaking). We used 50-ms non-overlapping windows spanning -500 to 1000 ms relative to each phrase and each patient. Following previous research (Chavez et al., 2025), for each window, we extracted spike counts and fit Poisson generalized linear models (GLM) with ridge regularization to predict neural responses from semantic features. Predictors included the first 100 principal components of mBERT embeddings (z-scored), phrase duration (z-scored), and their interactions, yielding 201 features total. Model performance was quantified as ΔLLH, the difference between the actual model log-likelihood and the median log-likelihood from 100 null models in which phrase-neural correspondence was shuffled. The analysis revealed that semantic encoding peaked ∼200-250 ms after visual onset and ∼300-400 ms before speech onset, indicating that hippocampal neurons encode semantic content during both retrieval and pre-articulatory preparation. Although our timing analysis (based on Patient 2 and Patient 3) suggested peaks at 300-400 ms before speech onset, we adopted the more conservative 200 ms pre-shift established by Chavez et al. (2025) using a larger cohort (n = 10 patients) to ensure robustness across patients.

### Electrode visualization for Figure 1A

Electrode visualization was performed using MNE-Python (version 0.24.0) with FreeSurfer’s fsaverage template brain. The visualization pipeline created an ultra-transparent glass brain (3% opacity) with highlighted hippocampal structures to clearly display electrode positions within deep brain regions. For each patient, DICOM images of the preoperative T1 anatomical MRI and the postoperative Stealth CT scans were acquired and converted to NIfTI format (Li et al., 2016). The CT was aligned to MRI space using FSL (M. Jenkinson & Smith, 2001; Mark Jenkinson et al., 2002). The resulting co-registered CT was loaded into BioImage Suite (version 3.5β1; (Joshi et al., 2011)), and the electrode contacts were manually localized. Electrodes’ coordinates were converted to the patient’s native space using iELVis MATLAB functions (Groppe et al., 2017) and plotted on the Freesurfer (version 7.4.1) reconstructed brain surface (Dale et al., 1999). Microelectrode coordinates are taken from the first (deepest) macro contact on the Ad-Tech Behnke Fried depth electrodes. RAVE (Magnotti et al., 2020) was used to transform each patient’s brain and electrode coordinates into MNI152 average space. The coordinates were plotted together on a glass brain with the hippocampus segmentation.

### Methods for cross-language neurons identification (Figure 2)

For each dataset group, we computed Pearson correlations between English and Spanish firing-rate responses across all matched words for each neuron. Neurons with zero variance in either language were excluded from inference and carried as NaN. For every remaining neuron, we assessed significance with a permutation test. Specifically, we generated a null distribution by randomly shuffling the Spanish word indices 1000 times and recomputing the correlation for each shuffle. The per-neuron p-value was the proportion of null correlations whose absolute value was greater than or equal to the absolute observed correlation. Neurons with p < 0.05 were classified as significant and then split by sign into significant positive (cross-language similar) and significant negative (cross-language anticorrelated). We set the random seed at the start of the analysis to ensure reproducibility.

At the population level, we summarized each group with two complementary tests. First, to ask whether the number of significant positive (and, separately, significant negative) neurons exceeded what would be expected from the per-neuron test’s false-positive rate, we used right-tailed binomial tests with chance probability p₀ = 0.025. This value corresponds to the expected false-positive rate from the two-tailed per-neuron permutation test (α = 0.05), in which only one tail represents positive correlations. The proportion of significant neurons in each dataset was visualized using pie charts (Figure 2), which displayed the percentage of neurons with significant positive cross-language correlations (purple), the expected proportion by chance (grey), and the remaining non-significant neurons (white).

We performed the same analysis at each layer of mBERT (see **Table S5**).

### Word-level contribution analysis for cross-language neurons (Table S1 and Table S2)

For each cross-language neuron in Study 1, across all stimulus sets, we decomposed the cross-language Pearson correlation into per-word contributions and defined “driver words” as the minimal set accounting for 80% of the positive contribution mass. We compared across-language neurons to size-matched bootstrap samples of non-across-language neurons (100 iterations per group) on three properties of these driver words: (1) selectivity (the percentage of the driver words), (2) semantic clustering (mean pairwise cosine similarity of driver word mBERT embeddings, tested against permutation null, permutation=1000, we named it as cluster index), and (3) effective dimensionality, the participation ratio (PR) of the driver words’ embedding matrix. For each neuron, we extracted the mBERT embeddings (Layer 11) of its driver words, performed PCA via singular value decomposition, and computed PR = (Σλ_i)² / Σλ_i², where λ_i are the eigenvalues. PR ranges from 1, when all variance is concentrated along a single principal axis, to the number of non-zero eigenvalues, when variance is uniformly distributed across all axes. It thus provides a continuous measure of how many independent semantic dimensions the driver words effectively span. A lower PR indicates that the driver words cluster along fewer embedding dimensions (i.e., occupy a more semantically constrained subspace), whereas a higher PR indicates that they spread across a broader range of semantic directions. To ensure a fair comparison, PR values of cross-language neurons were compared against size-matched bootstrap samples of non-cross-language neurons (100 iterations per group), so that any difference in participation ratio reflects genuine differences in the semantic geometry of driver words rather than differences in sample size.

In the read aloud task, we performed the same driver word analysis as in Study 1 (listening). Because the majority of results were reported at the word level, we did not extend this analysis to the phrase level. Because cross-language neurons vastly outnumbered non-cross-language neurons (61 vs. 12), the bootstrap procedure was inverted: in each of 100 iterations, 12 cross-language neurons were randomly subsampled (without replacement) and compared to the fixed set of 12 non-cross-language neurons.

### Whether there are language-specific neurons (Table S4)

To test whether individual hippocampal neurons were driven by the same words across languages or by different words in each language (language-specific or “separate” firing), we performed a word-level overlap analysis. For each neuron, firing rates were z-scored within each language across all words. A neuron was considered responsive to a word if its firing rate exceeded a fixed threshold (z > 1, that is, ∼top 16% responsive words). This yielded two binary response sets per neuron: English-responsive words and Spanish-responsive words. Responsive words were partitioned into three mutually exclusive categories: words eliciting responses in both languages (shared), words eliciting responses only in English, and words eliciting responses only in Spanish. To assess whether the observed overlap between English- and Spanish-responsive word sets exceeded chance expectations given each neuron’s marginal response rates, we used an exact hypergeometric test (without replacement). For each neuron, the null distribution assumed random overlap between English-responsive and Spanish-responsive word sets, conditioned on the total number of words, the number of English-responsive words, and the number of Spanish-responsive words. We computed the probability of observing an overlap greater than or equal to the empirical overlap under this null. Neurons with p < 0.05 were classified as showing overlap enrichment beyond chance. Based on this analysis, neurons were categorized as *pure-shared* (only shared responses), *pure-separate* (only language-specific responses), or *mixed* (both shared and language-specific responses).

### Contextual embedding extraction from multilingual BERT (Figure 3)

We extracted contextual word embeddings from multilingual BERT (mBERT) for complete English and Spanish versions of the transcripts for both stimulus sets, processing each language’s full stimulus independently before identifying matched word pairs for cross-linguistic analysis. The embedding extraction process was designed to maintain the contextual nature of transformer representations while ensuring that each word received naturalistic linguistic context from its own language stream. For each sentence in the transcripts (282 sentences in *set K*, 296 sentences in *set A*), we tokenized the text using mBERT’s tokenizer (bert-base-multilingual-cased) without adding special tokens initially, creating a flat sequential representation of all tokens across the entire stimuli. This approach allowed us to maintain accurate tracking of token positions and their relationships to original words in the transcript.

To preserve contextual information is crucial for mBERT’s contextualized representations, we implemented a sliding window strategy that provided each sentence with up to 509 tokens of preceding context (Zada et al., 2025). For each sentence, we calculated the available context window by subtracting the sentence length from the maximum allowable tokens (512 tokens minus 3 special tokens: [CLS], [SEP], [SEP]). The input sequence was then constructed following the format [CLS] + context_tokens + [SEP] + sentence_tokens + [SEP], where context tokens were drawn from immediately preceding text up to the calculated limit. This ensured that each word received rich contextual information from surrounding discourse while staying within mBERT’s architectural constraints.

The model processed each constructed input sequence to generate hidden state representations across all 13 layers (layers 0-12), with each layer capturing increasingly abstract linguistic features. We extracted embeddings specifically for the sentence tokens, excluding the context tokens and special tokens, by indexing into the appropriate positions of the output tensors. For words that were split into multiple subword tokens by the tokenizer, we averaged the embeddings across all constituent tokens to obtain a single word-level representation, ensuring that our analysis operated at the word level despite the subword tokenization scheme. This averaging preserved the dimensional structure (768 dimensions) while consolidating information from word pieces that collectively represent single lexical units.

The extraction process was performed identically for both English and Spanish transcripts, yielding parallel sets of contextualized embeddings where each word in the English version had a corresponding translation-equivalent word in the Spanish version. For each of the three podcast episodes and both languages, this resulted in a three-dimensional tensor of shape (n_layers × n_words × 768), where n_layers = 13 and n_words varied by stimuli set length.

These layer-wise embeddings captured the hierarchical processing of linguistic information, from surface-level features in early layers to more abstract semantic representations in deeper layers, enabling our subsequent analysis of cross-linguistic alignment patterns. The identical processing pipeline for both languages ensured that any observed differences in representation patterns could be attributed to linguistic rather than methodological factors.

### mBERT embeddings extraction for short phrase in read aloud task

We extracted mBERT embeddings using the pretrained *bert-base-multilingual-cased* model from HuggingFace Transformers. Each phrase was tokenized and passed through the model independently (i.e., without cross-phrase context), with input formatted as [CLS] phrase [SEP]. We extracted hidden states from Layer 11 (of 13 transformer layers), as middle-to-late layers have been shown to capture more semantic rather than syntactic information. For phrases containing multiple words, we obtained word-level embeddings by averaging the hidden states of all subword tokens belonging to each word. The resulting embeddings were 768-dimensional vectors for each word.

### Computing similarity between English-Spanish word embeddings in all layers

To investigate how multilingual language models represent semantic equivalences across languages, we analyzed the similarity between English and Spanish word embeddings extracted from parallel translated content. We obtained contextual embeddings for matched word pairs across three podcast episodes in both English and Spanish versions, extracting representations from multiple transformer-based models: multilingual BERT (mBERT) (Devlin et al., 2019; Pires et al., 2019) with 13 layers (layers 0-12). For each word pair, embeddings were extracted from every layer of these models to capture the hierarchical development of cross-linguistic representations.

For each stimulus set (Set A (audiobook: “*eat, pray, love*”) and Set K (*kurzgesagt*)), we computed Pearson correlations between the embedding vectors of each English-Spanish word pair (e.g., *“human-humano”*), yielding a correlation coefficient for every matched word across all layers. This approach quantified how similarly the models represent semantically equivalent words in different languages at various stages of processing depth.

To control for spurious similarities that might arise from random initialization or architectural properties rather than learned linguistic knowledge, we established a baseline using untrained versions of the same models with identical architectures but randomized weights. These baseline models underwent the same embedding extraction and correlation computation procedures as their trained counterparts, maintaining near-zero correlations across all layers as expected from random representations.

We aggregated correlations across all word pairs by computing means and standard errors for each layer, with error propagation applied when calculating the relative similarities to account for uncertainty in both trained and baseline measurements properly. The resulting layer-wise profiles revealed how cross-linguistic semantic alignment emerges and evolves through the depth of each model, with both stimulus sets showing progressive increases in correlation through the middle layers before reaching maximum alignment at Layer 11 (Figure 3B).

### Single neuron’s tuning vector analyses (Figure 3)

For each dataset (K, A1, A2, A3, KN-Neuropixel, read aloud task, conversation-listening, conversation-speaking), we concatenated single-unit firing-rate matrices across all words to form word × neuron matrices separately for English and Spanish. We extracted mBERT token embeddings for the matched words in each language (Layer 11, see above), then concatenated English and Spanish embeddings and ran a single PCA on the combined matrix to obtain a shared embedding basis. The resulting PCA scores were split back into English and Spanish, yielding a common set of semantic predictors with identical axes across languages (100 components).

Then, for every neuron, we fit two ridge regressions that map shared PCA embeddings to firing rates, one in English and one in Spanish. The ridge penalty was selected per neuron by cross-validation, using a grid of penalties and 5-fold CV jointly across languages (the chosen penalty minimized the average normalized prediction error over English and Spanish folds). With the selected penalty, we refit the English and Spanish ridge models and extracted, for each neuron, we got a *tuning vector*, that is, the vector of regression coefficients on the shared PCA semantic features. Each tuning vector (length = number of PCA components) specifies how strongly that neuron weights each semantic dimension to predict its firing rate.

We quantified neuron-wise similarity by correlating, for each neuron, its English tuning vector with its Spanish tuning vector (over PCA dimensions). To test whether English-Spanish similarity exceeded chance, we generated a null distribution by randomly permuting the Spanish word indices (1000 permutations, this procedure is same as we did in Figure 2), refitting the Spanish ridge model for every neuron with the same cross-validated penalty, and recomputing neuron-wise correlations. Per-metric p-values were computed as the proportion of null correlations whose absolute value was greater than or equal to the observed value (two-sided α = 0.05 test). Same as the method we used for Figure 2 (identify the cross-language neurons), we used right-tailed binomial tests with chance probability p₀ = 0.025 (corresponding to one tail of the per-neuron two-sided α = 0.05 test).

To evaluate whether the observed cross-language tuning similarity could be explained by within-language variability, we quantified the split-half reliability of tuning vectors for each neuron within English and within Spanish. For every iteration, word indices were randomly split into two equal halves, and independent ridge models were fit on each half using the neuron’s cross-validated penalty. The two resulting tuning vectors were correlated to obtain a within-language reliability value per neuron. This procedure was repeated 1000 times, generating a distribution of split-half correlations for each neuron and, when pooled across all neurons and both languages, a global distribution of within-language reliability. The vertical red line in Figure 3 (last panel) marks the mean cross-language tuning similarity across all neurons, shown relative to this within-language reliability distribution. We ran a paired Wilcoxon signed-rank test across neurons, testing whether per-neuron mean within-language reliability exceeded cross-language similarity.

### Constructing embedding dissimilarity matrix and neural dissimilarity matrix (Figure 4)

For both neural and semantic spaces, we computed *Representational Dissimilarity Matrices* (RDMs; (Diedrichsen & Kriegeskorte, 2017; Kriegeskorte & Kievit, 2013; Nili et al., 2014)) using cosine distance as the dissimilarity metric. The cosine distance between two vectors **x** and **y** is defined as:

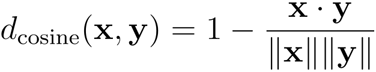

For a dataset with *n* words, this yields a condensed distance vector of length n*(n-1)/2 containing all unique pairwise distances. The neural RDM for language *l* was computed as:

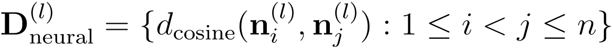

Where 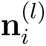 represents the neural population vector for word *i* in language *l*. Similarly, the semantic RDM was computed as:

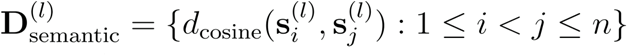

Where 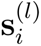 represents the mBERT embedding vector for word *i* in language *l*.

### Neural-semantic correspondence quantification

The correspondence between neural and semantic representational geometries was quantified by computing Spearman correlation between the vectorized RDMs (i.e., 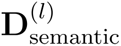 and 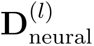). This correlation coefficient quantifies the degree to which neural dissimilarity patterns mirror semantic dissimilarity patterns. A high positive correlation indicates that words with similar meanings (small semantic distance) tend to evoke similar neural responses (small neural distance), supporting the hypothesis of semantic encoding in the neural population.

### Cross-language semantic map similarity analysis (Figure5)

We performed a cross-language similarity analysis comparing neural response patterns for matched English-Spanish word pairs to investigate the correspondence between neural representations of semantic content across languages. This analysis quantifies whether words that elicit similar neural responses in one language also elicit similar responses in another, providing evidence for shared or distinct neural geometry for semantic representation across languages.

Neural similarity between word representations was quantified using cosine distance, which measures the angular separation between neural population vectors independent of their magnitude. For each language, we computed pairwise cosine distances between all word pairs, this yielded similarity matrices *S*^(*E*)^ and *S*^(*S*)^ for English and Spanish, respectively, with values ranging from 0 (identical patterns) to 1 (maximally dissimilar patterns).

To assess the preservation of similarity structure across languages, we extracted the upper triangular portions of both similarity matrices (excluding the diagonal), yielding vectors of unique pairwise similarities. Then we use the Spearman correlation (Nili et al., 2014) to quantify the cross-language correspondence. A high positive correlation indicates that word pairs with similar neural representations in English tend to have similar representations in Spanish, suggesting a shared neural geometry for organizing semantic content across languages. To evaluate whether the observed cross-language correlation exceeded chance, we generated a null distribution by randomly shuffling the Spanish word labels 1000 times and recomputing the correlation after each shuffle. This procedure destroys the correspondence between matched English-Spanish word pairs while preserving each language’s internal structure. The resulting distribution of shuffled correlations represents the range of values expected if there were no true relationship across languages. The *p*-value was then calculated as the proportion of shuffled correlations whose absolute value was equal to or greater than the observed correlation (two-sided test, α = 0.05)

### Multidimensional Scaling (MDS) network visualization

In Figure 4B and Figure 4D, we employed classical Multidimensional Scaling (MDS) (Mead, 1992) combined with network visualization techniques to visualize the geometric organization of neural representations for matched word pairs across languages. This approach reveals the topological structure of semantic representations in neural space by projecting high-dimensional neural patterns onto interpretable low-dimensional manifolds while preserving pairwise dissimilarities. The resulting 2D coordinates were visualized as network graphs where nodes represent words and weighted edges encode distance between words.

### Two-dimensional histogram visualization in Figure 5C and Figure 5E

We constructed a two-dimensional histogram with 60 × 60 bins spanning the range [0, 1] for both dimensions to visualize the joint distribution of similarity values across languages. The bin counts were log-transformed using:

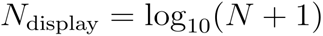

Where *N* represents the raw count in each bin, this transformation enhances visibility of the full density range while preventing singularities from empty bins.

### Local geometric cross-language prediction (Figure 5G-J)

To test whether preserved semantic geometry could support single-word neural decoding across languages, we implemented a local Procrustes alignment approach. All neural firing rate vectors were L2-normalized to unit length, which ensures that prediction accuracy reflects directional alignment in neural state space rather than differences in firing rate magnitude.

#### Anchor size optimization

We first determined the optimal number of anchor words by systematically varying *K* from 3 to 100. For each target word, we identified its k nearest semantic neighbors in English neural space (excluding the target itself), learned a local Procrustes rotation from these anchor pairs, and evaluated prediction accuracy on the held-out target.

#### Held-out prediction

For each target word with English response vector **e**_target_, we identified its nearest *K* semantic neighbors (*K* was the optimal anchor size from above analysis step) in English neural space using cosine distance (explicitly excluding the target word itself), then extracted these neighbors’ responses in both languages to form matrices **X***_E_* (English, *n_neurons_* × *k*) and **X***_S_* (Spanish, *n_neurons_* × *k*). After centering both matrices by subtracting column means (***μ****_E_* and ***μ****_S_*), we computed the cross-covariance matrix **M** = (**X***_S_* – ***μ****_S_*) (**X***_E_* – ***μ****_E_*)*^T^* and performed singular value decomposition **M** = **U∑V***^T^*. The optimal orthogonal rotation matrix was obtained as **Q** = **UV***^T^*, and the predicted Spanish response was computed as 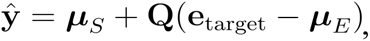, then L2-normalized to unit length. Prediction accuracy was quantified as cosine similarity and angular error 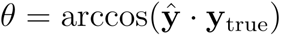 in degrees (arccos of cosine similarity).

#### Statistical significance

We tested predictions for all words in each dataset (datasets with matched English-Spanish words, including all stimulus sets in Study 1 and words from short phrases in Study 2). Statistical significance was assessed against 1000 null permutations where we (1) randomly shuffled English-Spanish word pairings using derangement (ensuring no correct translations), (2) randomly selected Spanish prediction targets, and (3) used randomly selected words (excluding the target) rather than true nearest neighbors as anchors for learning rotation matrix Q, thereby destroying both semantic correspondence and neighborhood structure while preserving neural response distributions. P-values were computed as the proportion of null means achieving equal or greater accuracy than the observed mean.

#### Sliding window analysis

To test whether the cross-linguistic rotation is local or global, we applied a sliding window approach. Instead of using the fixed nearest neighbors (positions 1 to *K*, where *K* represents the optimal anchor size), we shifted the window to use progressively more distant neighbors (for example, if *K*=17, then we have positions 2-18, 3-19, …, up to positions 84-100). If the rotation were strictly local, prediction accuracy should decay sharply beyond the immediate neighborhood.

#### Per-word trajectory analysis

To determine whether accuracy decay was gradual or sharp at the individual word level, we computed sliding window trajectories for each word separately. For each word, we calculated: (1) the overall slope of accuracy versus window position, and (2) the maximum single-step drop across adjacent windows. Sharp drops were defined as single-step decreases exceeding 2 standard deviations below the mean derivative across all words and windows.

#### Local geometry features

To examine what properties of local neighborhoods predict accuracy, we computed two metrics for each target word at optimal K (neighbors at positions 1 to k, excluding the target itself): (1) neighbor-cluster tightness, defined as the mean pairwise neural cosine similarity among the K neighbors. (2) The effective dimensionality of each anchor set using the participation ratio from PCA on the centered anchor matrix. The effective dimensionality reflects the neural geometric structure of the local neighborhood: low values indicate redundant anchors that span a low-dimensional neural subspace, while high values indicate structured anchors that span multiple dimensions. This metric allowed us to test the hypothesis that highly structured neighborhoods might predict better than redundant ones.

### Cross-language subspace alignment analysis (Figure 6C-D)

To examine how English and Spanish neural population representations relate to each other in geometric structure, we performed a cross-language subspace alignment analysis based on representational dissimilarity matrix (RDM, by cosine distance) and eigendecomposition. To enable eigendecomposition, we first converted cosine distances to a similarity matrix via 1 minus the RDM (where the subtraction is elementwise) to get the RSM (representation similarity matrix). We then applied double-centering (kernel centering) to remove the mean structure and obtain a centered Gram matrix (Borg & Groenen, 2005; Hofmann et al., 2008) directly in MATLAB (Statistics and Machine Learning Toolbox).

We then performed the eigendecomposition. Eigenvalues were sorted in descending order, and the corresponding eigenvectors were arranged accordingly. The eigenvectors define the principal semantic axes (or we called the “semantic readout axes” in *Result* sections) in the high-dimensional word space along which the neural population responses vary most strongly across words. That is, they are the directions that maximize variance in the centered word-word similarity structure.

To quantify representational dimensionality, we used the cumulative variance threshold, defined as the smallest number of components capturing at least 95% of the total variance. The top-k eigenvectors from each language formed orthonormal bases **U***_E_* and **U***_S_* representing the English and Spanish *semantic subspaces*, respectively. To quantify their geometric relationship, we adopted the alignment index from Elsayed et al. (2016)(Elsayed et al., 2016). We computed the variance-capture alignment index, which measures how much variance in one dataset’s representational structure is captured by another dataset’s subspace, normalized by the variance that the dataset could optimally capture on its own. The directional alignment is defined as:

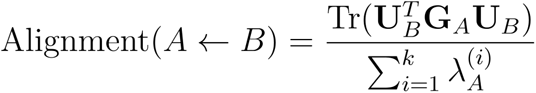

where **G***_A_* denotes the centered cosine Gram matrix for language A, which is mathematically equivalent to the second-moment (covariance-like) structure of neural activity in the cosine similarity space. *Tr*(.). is the matrix trace. Conceptually, **G***_A_* plays the same role as a covariance matrix, it captures how population responses co-vary across stimuli, which is computed from cosine similarities. **U***_B_* is the matrix of source language B’s top-k eigenvectors, and denominator is the sum of language A’s top-k eigenvalues. This metric ranges from 0 (no overlap) to 1 (perfect alignment), measuring what fraction of A’s top-k variance can be explained by projecting onto B’s top-k subspace. We computed bidirectional alignments Alignment (Eng ← Spa) and Alignment (Spa ← Eng), and reported their mean as a symmetric measure of overall subspace overlap between English and Spanish.

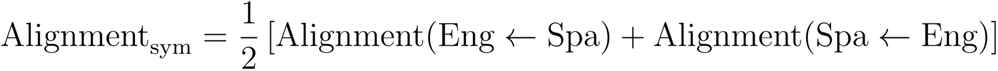

#### Permutation test

To test whether cross-language alignment exceeded chance, we performed 1000 permutation. In each permutation, the word order of the Spanish Gram matrix was randomly shuffled, disrupting the one-to-one correspondence between matched English-Spanish word pairs while preserving each language’s internal structure. For each shuffled instance, we recomputed the symmetric alignment index, producing a null distribution of alignment scores expected by chance. The empirical alignment was compared to this null distribution, and a permutation *p*-value was then calculated as the proportion of shuffled alignment whose absolute value was equal to or greater than the observed alignment (two-sided test, α = 0.05)

#### Compare to within-language subspace alignment

To assess the reliability of the across-language alignment, we computed within-language subspace alignment as a control. Because different subsets of words can produce distinct cosine-distance structures, splitting the words would alter the geometry of the representational dissimilarity matrix and thus would not reflect the reproducibility of the underlying subspace. Instead, we randomly divided the neurons into two equal halves (50% each) and repeated the procedure 1000 times. For each half, we recomputed the centered cosine Gram matrix, performed eigendecomposition, and re-estimated subspace alignment using the same alignment index calculation. To statistically evaluate whether the across-language alignment was significantly lower than within-language alignment, we compared the observed cross-language alignment value against the distribution of within-language alignments pooled across English and Spanish halves. We computed a *z*-score as the deviation of the observed value from the within-language mean, normalized by the within-language standard deviation. Significance thresholds followed standard criteria: |*z*| > 1.64 (*p* < 0.10), |*z*| > 1.96 (*p* < 0.05), and |*z*| > 2.58 (*p* < 0.01).

### Neuron-space coordination structure analysis (Figure 6E)

To determine whether English and Spanish engage the same neural ensemble with similar coordination patterns, we analyzed the covariance structure across neurons for each language. This analysis addresses whether cross-language semantic similarity arises from different neural subsets forming language-specific representations, or the same neurons reorganizing through different linear readouts.

#### Constructing neuron by neuron covariance matrix

For each language, we computed how pairs of neurons covary in their firing rates across all words. We first centered each neuron’s responses by subtracting its mean firing rate across words, then computed the covariance between all neuron pairs. This neuron-by-neuron covariance matrix quantifies patterns of coordinated neural activity: neurons with high positive covariance tend to increase their firing rates together across words, while neurons with negative covariance show opposite response patterns. We then performed eigendecomposition of each neuron covariance matrix to identify the dominant modes of coordinated neural activity. Each eigenvector represents a pattern of how neurons covary together, that is, neurons with high loadings on the same eigenvector tend to increase or decrease their firing rates together across words. The eigenvalues reflect the variance explained by each coordination mode, sorted from largest to smallest. We retained eigenvectors capturing greater than 95% of total variance. The number of eigenvectors required to reach this threshold was determined separately for each language. For cross-language comparisons, we used the minimum number of eigenvectors needed across both languages to ensure comparable dimensionality. We also quantified effective dimensionality using the participation ratio, which measures how many coordination modes contribute meaningfully to neural activity patterns. Then we quantified whether English and Spanish recruit the same patterns of neural coordination using the same alignment index for semantic subspaces alignment analyses.

#### Per-neuron participation strength

Also, to quantify each neuron’s contribution to the main coordination subspace, we computed participation strength as the sum of squared loadings across all retained eigenvectors. This metric reflects how strongly each neuron participates in the dominant coordination patterns. We computed the Spearman correlation between English and Spanish participation strengths to test whether the same neurons serve as the similar role of coordination in both languages.

#### Statistical validation via neuron-identity permutation

To test whether cross-language neural coordination alignment exceeds chance expectations, we performed 1000 permutations by randomly shuffling neuron identities between languages. This permutation test addresses the null hypothesis that neuron correspondences between English and Spanish are arbitrary. For each permutation, we randomly reordered the neurons in the Spanish data, breaking the neuron-to-neuron correspondence between languages while preserving each language’s word-evoked activity patterns. Then we recomputed the Spanish neuron covariance matrix from the permuted data, performed eigendecomposition and extracted the top coordination modes. Finally, we computed the alignment index and participation strength correlation under permutation. The permutation *p*-value was then calculated as the proportion of shuffled alignment whose absolute value was equal to or greater than the observed alignment (two-sided test, α = 0.05). If the observed values significantly exceeding the permutation distribution indicate that English and Spanish specifically recruit the same neurons with matched coordination patterns.

## Acknowledgements

We thank Joshua Adkinson, Justin Fine, Victoria Gates, Suzanne Kemmer, George Kokalas, and Steven Piantadosi for their assistance.

