## Supplemental figures for "Shared neural geometries for bilingual semantic representations in human hippocampal neurons"

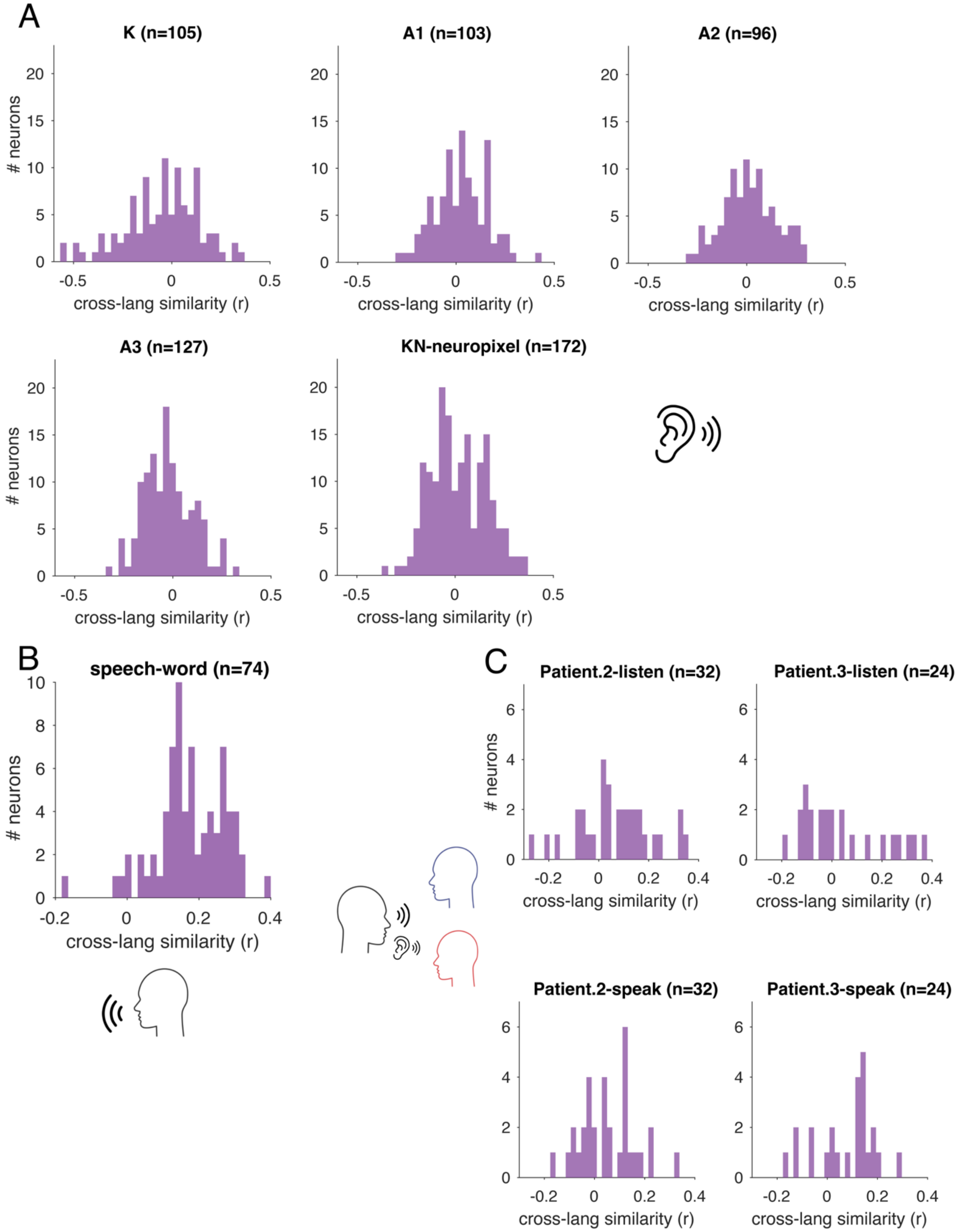


**Figure S1. The distribution of cross-language tuning similarity (Pearson correlation between English and Spanish tuning vectors) across all neurons for each study.**
Related to Figure 3.

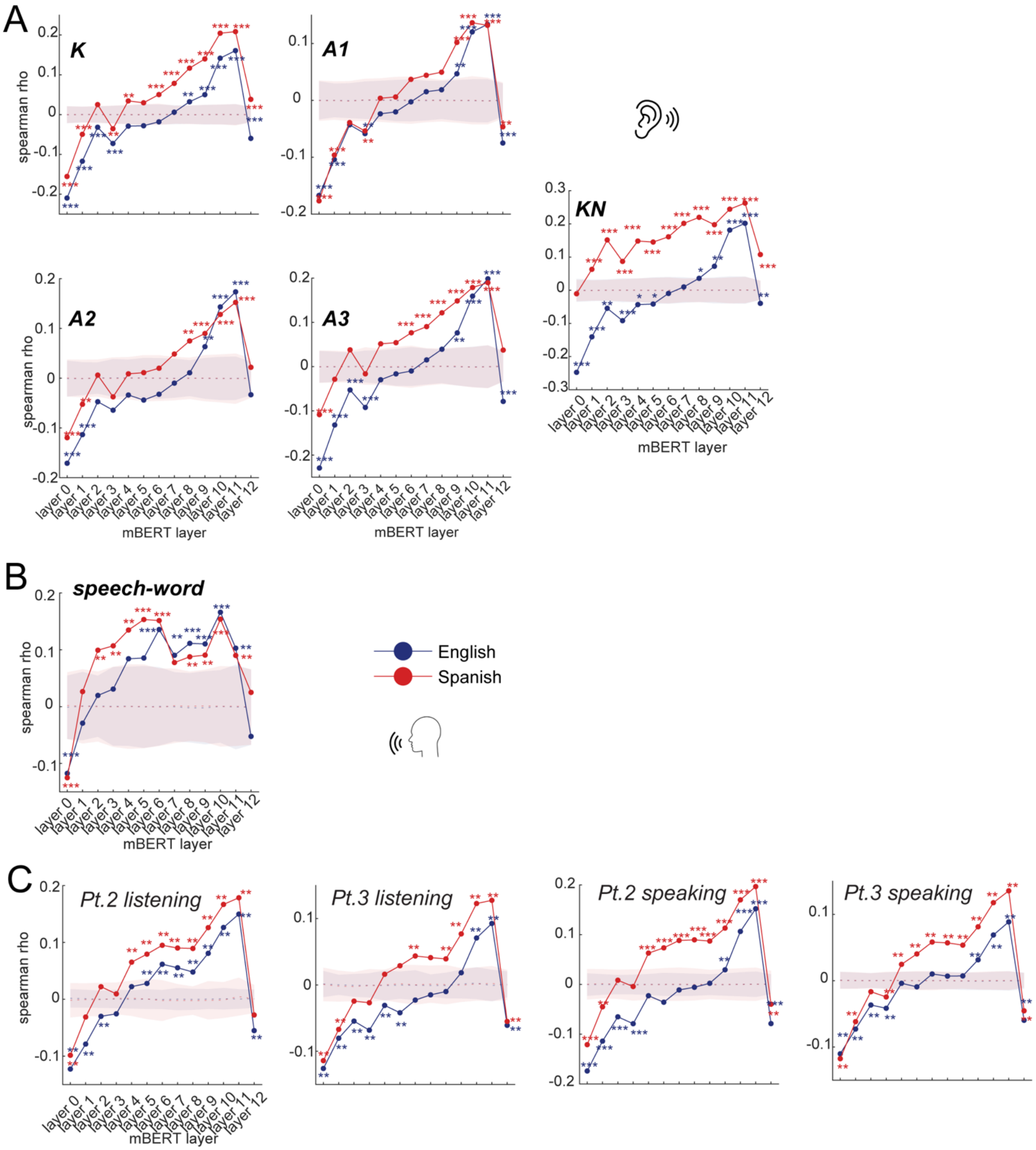


**Figure S2. Brain-mBERT alignment at each mBERT layer.**
We repeated the brain-mBERT alignment analyses (Figure 4) across all 13 mBERT layers (layers 0-12) for each stimulus set. Across all five stimulus sets (K, A1, A2, A3, and KN), speech production (word-level) as well as the conversation task, the correlation between neural and semantic RDMs was negative in layers 0-1, progressively shifted toward positive values around layers 4-6, and peaked with statistical significance in layers 9-11.
**p* < .05, ***p* < .01, ****p* < .001, permutation test (1000 iterations)
Related to Figure 4.

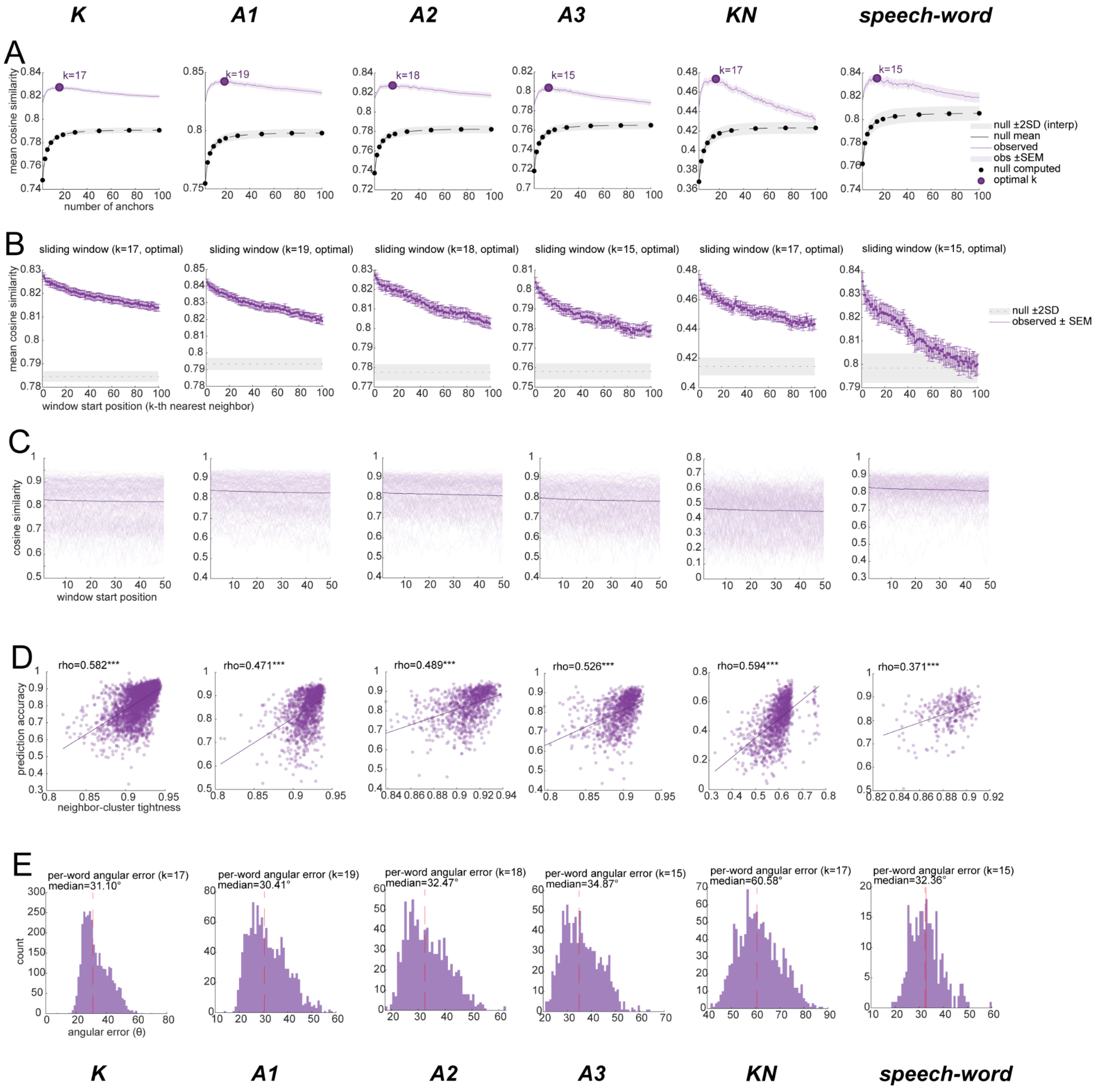


**Figure S3. Extended results for local geometric structure enables cross-language prediction** (A) Optimal anchor size. Mean cosine similarity between predicted and actual Spanish neural response vectors as a function of anchor set size (k = 3-100). Purple line shows observed prediction accuracy (±SEM); black dashed line shows null distribution mean with ±2 SD band (gray shading). Black dots indicate anchor sizes where null was explicitly computed (1000 permutations). Large purple dot marks optimal k for each dataset. Prediction accuracy peaks at k = 15-19 depending on dataset and remains stable across anchor sizes. The high null baseline reflects shared global covariance between the two neural spaces; the consistent observed advantage above null reflects the additional precision gained from true semantic correspondence. (B) Sliding window analysis. Mean cosine similarity as a function of window start position at optimal k. Position 1 uses neighbors ranked 1-k; position n uses neighbors ranked n to n+k-1. Accuracy decays by less than 1° across 100 window positions and remains far above null throughout, indicating the cross-linguistic transformation approximates a near-global rotation rather than a strictly local one.

(C) Per-word sliding window trajectories. Individual word trajectories across window positions. Near-horizontal profiles indicate gradual decay; absence of clustered sharp drops suggests no discrete neighborhood boundaries.

(D) Neighborhood tightness predicts accuracy. Relationship between neighbor-cluster tightness (mean pairwise neural cosine similarity among k anchors) and prediction accuracy for each word. Strong positive correlations (Spearman $\rho = 0.37$-$0.59$, all $p < 0.001$) indicate that words in tighter, more coherent neighborhoods are predicted more accurately.

(E) Per-word angular error distribution. Histograms of angular error ($\theta = \arccos(\text{cosine similarity})$) at optimal k (i.e., anchor size in panel A).

Columns: K = Kurzgesagt (3457 words); A1, A2, A3 = audio books sets 1-3; KN = Kurzgesagt-Neuropixel; speech-word = read-aloud task.

*$p < .05$, **$p < .01$, ***$p < .001$, permutation (1000 iterations)

Related to Figure 5.

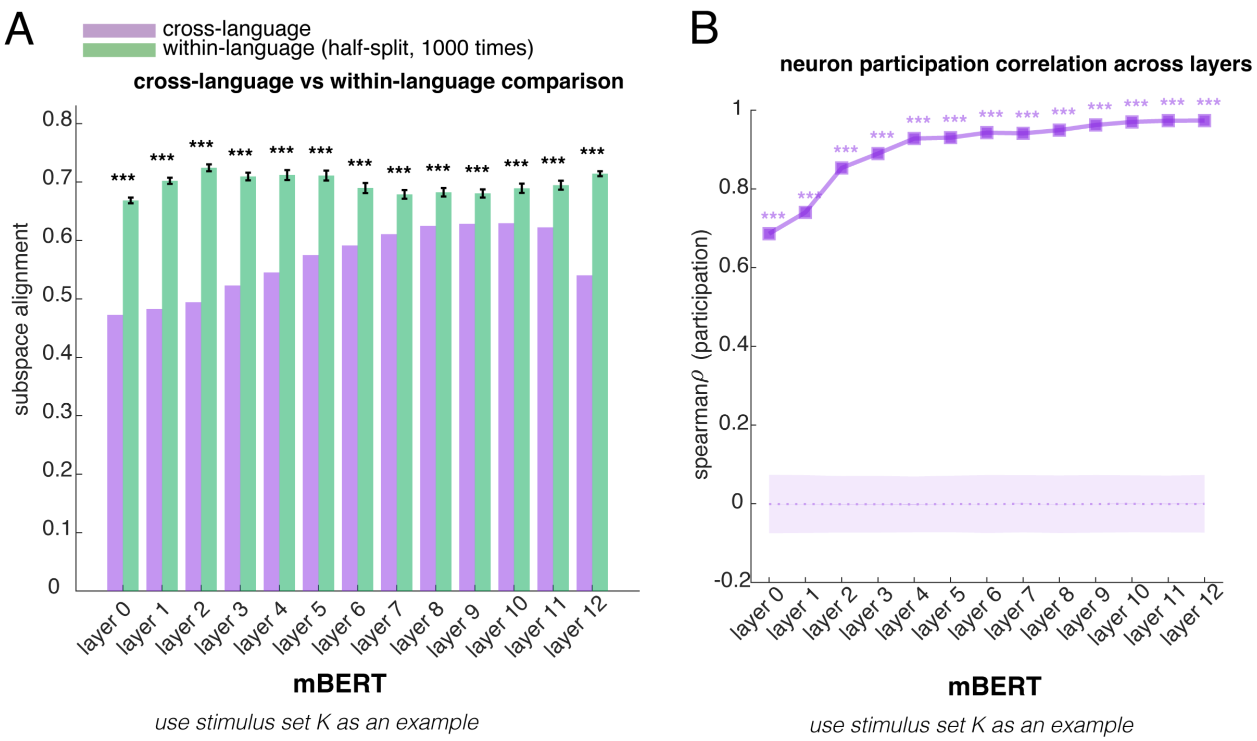


**Figure S4. Cross-language subspace alignment and "neuron" participation in subspace construction at every layer of mBERT.**

(A) Cross-language subspace alignment (using stimulus set K as an example) is significantly smaller than within-language half-split baselines at every layer of mBERT.

(B) If we treated the embedding dimension of mBERT as a neuron, the per-artificial neuron participation strength in the top subspace (explaining > 95 % variance) was highly correlated between languages (all rho>0.650, all p<0.001) at every layer of mBERT, showing that mBERT utilized the same "hardware" to construct language-specific semantic readout axes to achieve the shared meaning.

**p < .05, **p < .01, ***p < .001*, permutation test (1000 iterations)

Related to Figure 6.

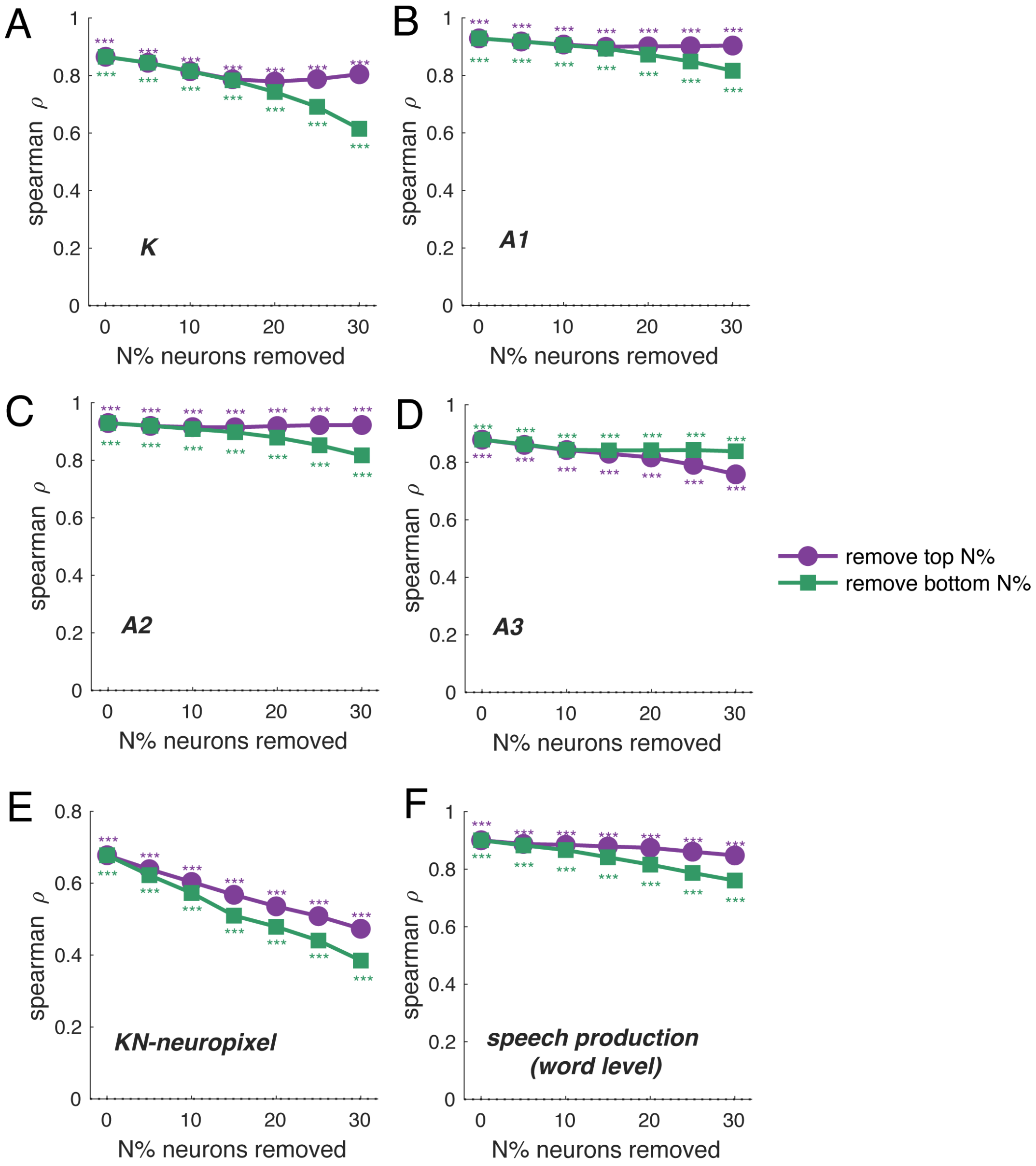


**Figure S5. The robustness of per-neuron participation strength.**
(A-E) We progressively removed the top 5-30% highest-participation neurons and recomputed the Spearman correlation at each step. The Spearman rank correlation remained very significant even after removing the top 30% of neurons (all stimulus sets, K, A1, A2, A3, KN-neuropixel in Study 1 and word speech in Study 2, $p<0.001$; purple line). We then progressively removed the bottom 5-30% lowest-participation neurons and recomputed the correlation at each step. The Spearman rank correlation remained very significant (all stimulus sets, K, A1, A2, A3, KN-neuropixel in Study 1, and word speech in Study 2, $p<0.001$; green line). Together, these results demonstrate that the effect is not driven by a small subset of extreme neurons.
$^{*}p < .05$, $^{**}p < .01$, $^{***}p < .001$, permutation (1000 iterations)
Related to Figure 6.

### SUPPLEMENTAL INFORMATION

**Table S1. Driver word properties of cross-language versus non-cross-language neurons (Study1).**

Cross-language neurons required significantly fewer words to explain their cross-language correlations, confirming that a particular subset of words disproportionately contributes. However, these driver words did not cluster into discernible semantic categories. Indeed, the mean semantic cluster index was identical between groups. That is, driver words of cross-language neurons were no more semantically similar to one another than those of non-cross-language neurons, which argues against the possibility that these neurons simply respond to a single named category (e.g., “animals” or “emotions”). Critically, despite equivalent pairwise semantic similarity, the driver words of cross-language neurons spanned fewer independent semantic dimensions. Because pairwise cosine similarity is a scalar average that collapses all directional information, it is insensitive to the anisotropy of the distribution; two sets of words can have identical average pairwise distances yet differ in how their variance is distributed across principal axes. Therefore, we calculated the participation ratio (PR), which provides a continuous measure of how many independent semantic dimensions the driver words effectively span. A lower PR indicates that the driver words cluster along fewer embedding dimensions (i.e., occupy a more semantically constrained subspace), whereas a higher PR indicates that they spread across a broader range of semantic directions. We found the driver words have lower participation ratio than non-cross-language neurons.

| **Stimulus** | ***n* (cross-language)** | **Selectivity (K80%)** | | | **Cluster Index *(similarity)*** | | | **Participation Ratio *(dimensionality)*** | | |
|---|---|---|---|---|---|---|---|---|---|---|
| | | **Cross-Language neurons** | **Non-cross-language neurons** | ***d*** | **Cross-Language neurons** | **Non-cross-language neurons** | ***d*** | **Cross-Language neurons** | **Non-cross-language neurons** | ***d*** |
| K | 6 | 21.6 ± 6.4 | 18.3 ± 5.9 | 0.56 | 0.663 ± 0.002 | 0.664 ± 0.003 | -0.092 | 66.5 ± 3.4 | 65.6 ± 3.4 | 0.27 |
| A1 | 20 | 16.3 ± 5.8 | 25.6 ± 14.1 | $-0.66^{**}$ | 0.666 ± 0.011 | 0.667 ± 0.004 | -0.295 | 55.3 ± 12.6 | 62.7 ± 6.5 | $-1.13^{***}$ |
| A2 | 11 | 19.2 ± 9.1 | 24.3 ± 14.6 | -0.35 | 0.696 ± 0.004 | 0.691 ± 0.005 | $0.927^{**}$ | 47.4 ± 4.4 | 47.9 ± 5.3 | -0.10 |
| A3 | 8 | 17.2 ± 11.1 | 30.0 ± 17.1 | $-0.75^{*}$ | 0.693 ± 0.019 | 0.690 ± 0.006 | 0.536 | 41.0 ± 17.4 | 49.7 ± 6.7 | $-1.28^{***}$ |
| KN-neuropixel | 9 | 26.6 ± 3.7 | 28.4 ± 5.2 | -0.35 | 0.651 ± 0.004 | 0.652 ± 0.004 | -0.294 | 56.9 ± 2.6 | 57.6 ± 3.4 | -0.19 |
| **Pooled** | **54** | **19.3 ± 8.0** | **25.7 ± 13.4** | **$-0.48^{***}$** | **0.673 ± 0.019** | **0.673 ± 0.015** | **0.017** | **53.4 ± 12.4** | **57.2 ± 8.7** | **$-0.44^{**}$** |

*Table Note.* Values in each cell are mean ± SD.
d = Cohen’s d (negative indicates cross-language neurons < non-cross-language neurons).
Selectivity (K80%): percentage of words needed to explain 80% of positive cross-language correlation mass.
Cluster Index: mean pairwise cosine similarity among driver words in mBERT embedding space.
Participation Ratio: effective dimensionality of driver word embeddings.
$^{*}p < .05$, $^{**}p < .01$, $^{***}p < .001$

Related to Figure 2.

**Table S2. Driver word properties of cross-language versus non-cross-language neurons (read aloud task, Study 2, word level).**

In the read aloud task, we performed the same driver word analysis as in Study 1 (listening). Because most results were reported at the word level, we did not extend this analysis to the phrase level. Of 73 valid neurons, 61 were classified as cross-language neurons (83.6%), leaving only 12 non-cross-language neurons.

Because cross-language neurons vastly outnumbered non-cross-language neurons (61 vs. 12), the bootstrap procedure was inverted: in each of 100 iterations, 12 cross-language neurons were randomly subsampled (without replacement) and compared to the fixed set of 12 non-cross-language neurons. Although the cluster index differed significantly between groups, the absolute difference was negligible (Δ = 0.010), suggesting that the semantic similarity structure of driver words was virtually identical across neuron types; the statistical significance likely reflects the large effective sample size in the bootstrap rather than a meaningful biological difference.

| **Stimulus** | ***n* (cross-language)** | **Selectivity (K80%)** | | | **Cluster Index *(similarity)*** | | | **Participation Ratio *(dimensionality)*** | | |
|---|---|---|---|---|---|---|---|---|---|---|
| | | **Cross-Language neurons** | **Non-cross-language neurons** | ***d*** | **Cross-Language neurons** | **Non-cross-language neurons** | ***d*** | **Cross-Language neurons** | **Non-cross-language neurons** | ***d*** |
| Word-speech | 61 | 17.4 ± 3.0 | 33.8 ± 16.4 | $-2.35^{***}$ | 0.807 ± 0.006 | 0.797 ± 0.004 | $1.722^{***}$ | 28.4 ± 4.0 | 34.7 ± 5.6 | $-1.47^{***}$ |

*Table Note.* Values in each cell are mean ± SD.
d = Cohen's d (negative indicates cross-language neurons < non-cross-language neurons).
Selectivity (K80%): percentage of words needed to explain 80% of positive cross-language correlation mass.
Cluster Index: mean pairwise cosine similarity among driver words in mBERT embedding space.
Participation Ratio: effective dimensionality of driver word embeddings.
$^{*}p < .05$, $^{**}p < .01$, $^{***}p < .001$.
Related to Figure 2.

**Table S3. Cross-language neurons did not affect shared representation geometry.**

We chose the Steiger's Z-test for dependent correlation difference significance test. To test whether cross-language neurons disproportionately contribute to cross-language representational alignment, we compared the observed drop in English-Spanish RDM (as well as in neural-LLM correlation) upon removing cross-language neurons against a null distribution generated by randomly excluding the same number of neurons 1000 times. The results show no differences (all p>0.400), indicating that cross-language neurons do not contribute more to cross-language geometric similarity than a random subset of equal size. We repeated such analysis for every stimulus set.

| neural-LLM | | K | A1 | A2 | A3 | KN |
|---|---|---|---|---|---|---|
| r_with | spanish | r=0.209*** | r=0.132*** | r=0.152*** | r=0.190*** | r=0.263*** |
| | english | r=0.161*** | r=0.134*** | r=0.173*** | r=0.199*** | r=0.202*** |
| r_without | spanish | r=0.208*** | r=0.164*** | r=0.153*** | r=0.188*** | r=0.259*** |
| | english | r=0.160*** | r=0.174*** | r=0.171*** | r=0.199*** | r=0.199*** |
| Steiger's Z (r_without vs. r_with) | spanish | z = 0.329, p = 0.742 | z=-21.751, p<0.001 | z=-0.563, p=0.573 | z = 0.846, p = 0.397 | z=2.105, p=0.035 |
| | english | z = 0.675, p = 0.500 | z=-28.278, p<0.001 | z=1.029, p=0.303 | z = 0.723, p = 0.469 | z=2.136, p=0.033 |
| Permutation p-value for (Δr vs. Δr_random) | spanish | p=0.751 | p=1.00 | p=0.764 | p=0.511 | p=0.206 |
| | english | p=0.747 | p=1.00 | p=0.636 | p=0.506 | p=0.041 |

| English-Spanish neural geometry similarities | K | A1 | A2 | A3 | KN |
|---|---|---|---|---|---|
| r_with | r=0.477*** | r=0.255*** | r=0.281*** | r=0.358*** | r=0.385*** |
| r_without | r=0.474*** | r=0.341*** | r=0.223*** | r=0.316*** | r=0.379*** |
| Steiger's Z (r_without vs. r_with) | Z = 3.758, p = 0.0001 | Z = -62.502, p<0.0001 | Z = 2.447, p = 0.01 | Z = 3.963, p < 0.0001 | Z = 3.656, p = 0.0002 |
| Permutation p-value for (Δr vs. Δr_random) | p=0.773 | p =1.000 | p=0.736 | p=0.486 | p=0.606 |

*Table Note.* r_with = Cross-language RDM similarity (with cross-language neurons);
r_without = Cross-language RDM similarity (without cross-language neurons);
Δr = r_with − r_without;

Δr_random = mean Δr from random neuron exclusion (of equal size) (mean ± SD)
$^{*}p < .05$, $^{**}p < .01$, $^{***}p < .001$
Related to Figure 4 and Figure 5.

**Table S4. No language-specific neurons**

In the stimulus set K and KN (Study1), at the single-neuron level, no neuron was selectively active in only one language. Every neuron exhibited a mixture of shared and language-specific word responses. In the stimulus sets A1, A2, A3 (Study 1) and read aloud task (speech production, word level, Study 2), although a small number of neurons appeared responsive in only one language, these counts did not exceed the rate expected by chance (binomial test, all $p > 0.300$). For all stimulus sets, across the population, cross-language overlap was significantly enriched relative to chance (Wilcoxon signed-rank test, all $p<0.001$; Cohen's $d > 0.30$). This matches our broader conclusion: shared meaning is not implemented by special neurons, but by population geometry.

| | **stimulus-K** | **stimulus-A1** | **stimulus-A2** | **stimulus-A3** | **stimulus-KN** | **Speech-word** |
|---|---|---|---|---|---|---|
| **mixed** | 105 | 101 | 92 | 119 | 172 | 72 |
| **Pure shared** | 0 | 0 | 0 | 0 | 0 | 0 |
| **Pure selective** | 0 | 2 | 4 | 8 | 0 | 2 |
| **selective-neurons above chance?** | p=1.000 | p=0.967 | p=0.713 | p=0.303 | p=1.000 | p=0.890 |
| **cross-language overlap exceeded chance** | p<0.001 | p<0.001 | p<0.001 | p<0.001 | p<0.001 | p<0.001 |

Related to Figure 2.

**Table S5. “Cross-language neurons” in mBERT (all layers)**

At the single-unit level: in mBERT, the vast majority of ‘units’ are cross-language correlated units. All units of higher layers of mBERT (>=layer 6) are cross-language correlated units

| Layer | K | A1 | A2 | A3 | KN | Speech-word |
|---|---|---|---|---|---|---|
| layer0 | 99.7% | 99.5% | 97.3% | 97.0% | 99.2% | 83.1% |
| layer1 | 99.9% | 100% | 99.2% | 100% | 99.9% | 90.4% |
| layer2 | 100% | 100% | 100% | 100% | 100% | 94.1% |
| layer3 | 100% | 100% | 100% | 100% | 100% | 97.8% |
| layer4 | 100% | 100% | 100% | 100% | 100% | 99.5% |
| layer5 | 100% | 100% | 100% | 100% | 100% | 99.7% |
| >=layer6 | 100% | 100% | 100% | 100% | 100% | 100% |

Related to Figure 2 and Figure 6.
